# Representational drift shows same-class acceleration in visual cortex and artificial neural networks

**DOI:** 10.1101/2025.11.05.686897

**Authors:** Jinke Liu, Amr Farahat, Martin Vinck

**Affiliations:** Ernst Strüngmann Institute for Neuroscience in Cooperation with Max Planck Society; Donders department of Neurophysics, Nijmegen University

**Keywords:** representational drift, dimensionality reduction, population coding

## Abstract

The neural code is not fixed but shows substantial representational drift over time. It has been proposed that representational drift reflects continuous learning resulting from input-dependent plasticity. Using theoretical analysis and simulations in artificial neural networks, we show that input-dependent plasticity entails a same-class-acceleration principle: Representational drift for a given class of stimuli is predominantly caused by presenting that class of stimuli, rather than presenting other classes of stimuli. We analyze electrophysiological recordings of mouse visual cortex to examine whether within-session representational drift is consistent with this principle. Within-session representational drift was not explained by changes in behavioral state. Instead, it showed a systematic temporal structure that reflected sensory experience rather than behavior. Drift for a given set of stimuli accelerated during blocks presenting that set of stimuli and slowed down during blocks in which other stimuli were presented. Thus, sensory inputs, not elapsed time, organize representational drift in a stimulus-specific manner, consistent with theoretical predictions for training neural networks.

## INTRODUCTION

Neural responses to the same stimulus can vary significantly across repetitions or days. Unlike noise, this trial-to-trial variability continuously accumulates and thus exhibits some form of persistence. Such gradual changes in neural representations are referred to as *“representational drift”* and have been observed in many cortical structures(Aschauer et al., 2019; Chen et al., 2015; Schoonover et al., 2021; Rokni et al., 2007; Low et al., 2020; Singh et al., 2019; Ziv et al., 2013; Buzsáki and Tingley, 2018; Keinath et al., 2022; Tang et al., 2025; Sotomayor-Gomez et al., 2023), and on different time scales, e.g. within a session on the same day (Cowley et al., 2020; Sotomayor-Gomez et al., 2023) or across multiple sessions (Deitch et al., 2021; Bauer et al., 2023). Representational drift poses an important problem, namely how the brain maintains stable behavior in light of continuously changing codes, but may also provide a window as to how neural representations change as a function of continuous learning (Micou and O’Leary, 2023; Zhong et al., 2025).

Although representational drift may in part reflect fluctuations in behavioral state or excitability (Low et al., 2020; Vinck et al., 2015; McGinley et al., 2015; Avitan and Stringer, 2022; Bimbard et al., 2023; Rule et al., 2019; Sadeh and Clopath, 2022; Micou and O’Leary, 2023; Delamare et al., 2024), it may also result from gradual changes in synaptic connectivity. These changes could result from random synaptic turnover that introduces stochastic variability over time (Attardo et al., 2015; Chambers and Rumpel, 2017; Raman and O’Leary, 2021). Representational drift may also reflect learning-related mechanisms or ongoing circuit reorganization in response to experience (Bauer et al., 2023; Rule and O’Leary, 2022; Driscoll et al., 2022; Zhong et al., 2025; Devalle et al., 2025).

Via theoretical analyses and simulations we show here that when neural networks are trained with stochastic gradient descent, they show a *same-class-acceleration* principle: If synaptic weights are updated due to the presentation of a stimulus of category *c*, then these weight changes produce larger changes in neural activity for *same-category* probes (or probes similar in representational space) than for *different-category* probes. Thus, we predict that in neural data, the speed of drift should not be constant but scale according to sensory experience, and that representational drift for a given set of stimuli accelerates when those stimuli are presented, and slows down when other stimuli are presented. By analyzing multi-areal electrophysiological recordings in mouse visual cortex, we indeed show that representational drift shows a characteristic *M-shaped* pattern of acceleration and deceleration across blocks that is consistent with the same-class-acceleration principle we derived theoretically.

## RESULTS

### Theoretical analyses

We consider neural networks trained via stochastic gradient descent (SGD). We will show that the experience-dependent changes in responses to a stimulus class *c* are expected to be large after presenting class *c*, but smaller during the experience of different stimulus classes. We derive this property theoretically for neural networks, and we substantiate it empirically by training neural networks and analyzing changes in neural representations from iteration to iteration. Specifically, our proposition is that when we present a stimulus from class *c*, leading to an update in the parameters of the neural network, the responses of the network show a stronger change for a same-class test probe of class *c* than a test probe of a different class.

Detailed theoretical proofs are given in the Methods and briefly summarized here. We first consider the case of a single-layer neural network, and then generalize to a multi-layer neural network. Suppose we train this network via SGD with a training step for an input *x* of class *c*, and then examine the change in the network representations, the logits 𝓏, for a test probe *x*′. We show that same-class test-probes show, on average, a larger change, i.e.

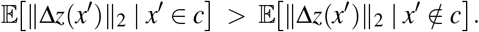

if within-class input features have a greater similarity than across-class input features, i.e.

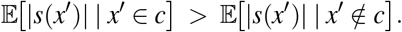

Here *s*(*x*_*c*_, *x*^′^ ) = ⟨*x*_*c*_, *x*′ ⟩ is the similarity of input features between the training and test probe. This inequality, as in similar cases below, is trivially satisfied if the train and test probe are identical, i.e. *x* = *x*_*c*_.

A similar behavior is expected in the last layer of a trained, multi-layer (i.e. deep) neural network, Let *a*^(*L*)^(*x*) be the activation in the last layer of the neural network and 𝓏 ^(*L*+1)^(*x*) = *W*^(*L*+1)^*a*^(*L*)^(*x*) ∈ℝ^*K*^ the logits. It can be shown that (see Methods) for any test probe *x*^*′*^, the change in its logits is

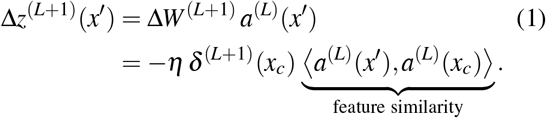

Taking norms,

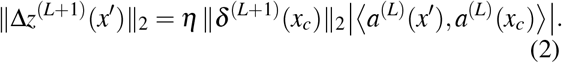

Equation (2) shows that, for fixed (*x*_*c*_, *c*), the probe dependence is entirely through the scalar similarity *s*(*x*^*′*^, *x*_*c*_) ≡ ⟨ *a*^(*L*)^(*x*^*′*^ ), *a*^(*L*)^(*x*_*c*_) ⟩ of last-layer features, measured in terms of neural network activations. Hence a condition for accelerated same-class representational drift is

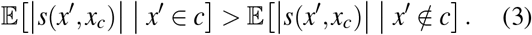

In trained neural networks, this condition is typically satisfied as within-class inputs cluster in feature space, which thus increases the inner product ⟨ *a*^(*L*)^(*x*^*′*^ ), *a*^(*L*)^(*x*_*c*_) ⟩ for *x*^′^ ∈*c*. This establishes same-class accelerated drift in the readout layer.

Finally, we consider the case of representations in any layer of a deep neural network. We derive a Cauchy-Schwarz bound on the changes in neural representations over time and show that this bound can be written as a product of forward and backward similarity in the neural network,

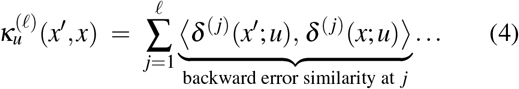

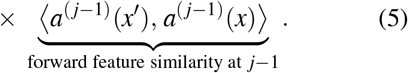

Thus, for pre-activations at any layer, representational drift depends on the (gated) forward × backward similarities. Since training increases both forward feature similarity and backward alignment within class—especially in deeper layers—this yields a theoretical explanation for same-class acceleration that becomes more pronounced with depth and training.

### Neural network simulations

To test these theoretical predictions, we first analyzed changes in a single-layer neural network performing classification on MNIST, trained via SGD. Changes in neural representations show within-class acceleration; e.g. presenting a stimulus from class 0 causes the strongest change in representations for a test probe from class 0. This pattern is consistently found for all classes (Figure 1).

**Figure 1.**
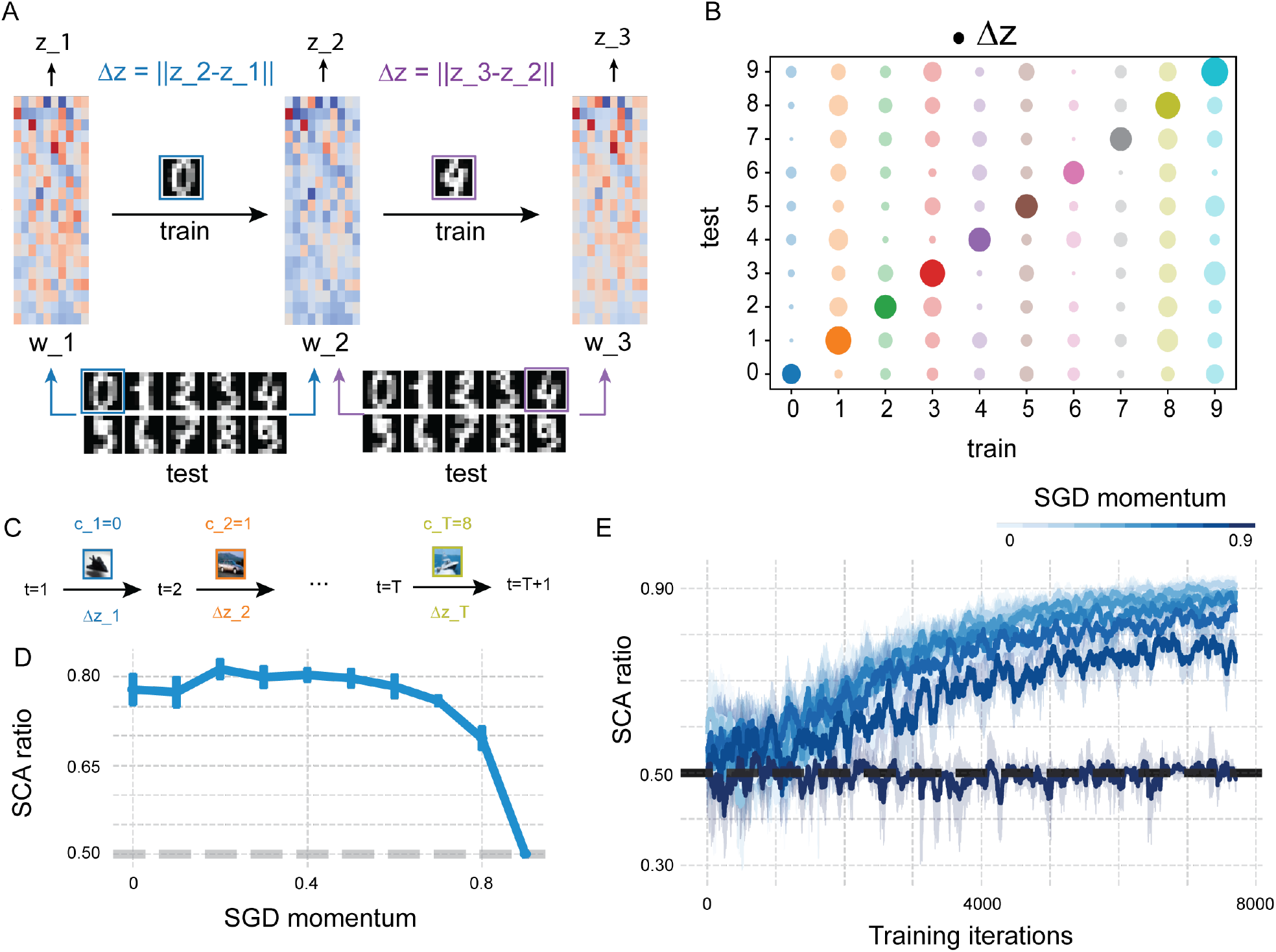
Neural network simulations. **(A)** Schematic of a logistic regression model trained on the MNIST digit classification task. **(B)** Changes in neural representations after training Δ**z**. The diagonal corresponds to test classes that match the training class. **(C)** Schematic of a convolutional neural network (CNN) trained on the CIFAR-10 classification task, each timestep a batch of the same class image was used to train the network. **(D)** Same-Class-Acceleration (SCA) ratio as a function of momentum after the network is fully trained. The ratio reflects the percentage of cases where the largest representational change occurs for classes matching the training class. Chance level is 0.5 and shown in dashed line. **(E)** Learning curves showing SCA ratio over training iterations for different momentum settings.

Next, we consider a multi-layer neural network, which we trained on 10 classes with a three-layered convolutional neural network. After each minibatch, in which only images from one class were presented, we evaluated the change in representations in the hidden states of the penultimate fully-connected layer for all stimuli of all classes in the test set. We computed the number of times that a given test class showed the largest representational change when the training class was of the same class, *N*_same_, and the number of times that the given test class showed the largest change when the training class was of another class, *N*_other_. We then computed a same-class-acceleration (SCA) ratio as *N*_same_*/*(*N*_same_ + *N*_other_). We found that this SCA-ratio was substantially higher than chance level (80% vs. 50%). We expected this effect to be greater when we trained with less momentum (in the SGD optimizer), as momentum causes effective memory across training batches. Indeed, as we increased the momentum hyperparameter, the SCA-ratio dropped to chance level. We furthermore tested the analytical prediction that the SCA-ratio should increase with training. Indeed, the SCA-ratio increased steadily with training, reaching values around 0.9 after approx. 7500 training iterations. This yields robust evidence for the existence of same-class-accelerated representational drift in an artificial network trained on object recognition, aligning with our analytical derivations.

### Neural representation of visual stimulus drifts across presentation blocks

We then analyzed electrophysiological and calcium imaging data. The main dataset analyzed consisted of extra-cellular, electrophysiological recordings made with Neuropixels (Allen Institute; Brain Observatory Paradigm 1.1, Siegle et al. (2021)). Sessions were divided into multiple blocks during which visual stimuli were presented. In each visual stimulation block, a different set of visual stimuli was presented, e.g. drifting gratings or natural movies. We initially analyzed a subset of sessions that contained blocks of drifting grating trials (the “Session A” protocol, Figure 2A). Drifting grating stimuli were presented in three different blocks that were separated by gaps that consisted primarily of natural images and videos. In the drifting-grating block, each drifting-grating stimulus was presented for two seconds with an inter-trial interval of one second. Each grating pattern could move in one of the eight different directions and was repeated for about 25 trials in random order. Firing rates were calculated for the first 200 ms after stimulus onset. We constructed Representational Dissimilarity Matrices (RDMs) copmuting the pairwise Euclidean distance between the normalized population vectors of all trials across the three blocks (Figure 2B). Consistent with the known tuning properties of the V1 neurons, low-dimensional t-SNE embeddings showed distinct clusters for the eight stimulus directions (Figure 2C,E). The example in Figure 2E suggests that a substantial fraction of the variability in the low-dimensional embedding space was driven by the block identity (i.e. 1, 2 or 3), rather than the orientation of the drifting grating. This variability was not accounted for by the average firing rates during the block (Figure 2D). The representational drift was accounted for by heterogeneous changes in neuronal responsiveness across blocks (e.g. see Figure 2C): The tuning curves of visual neurons were modulated in various additive and multiplicative ways (Figure S2, see Methods).

**Figure 2.**
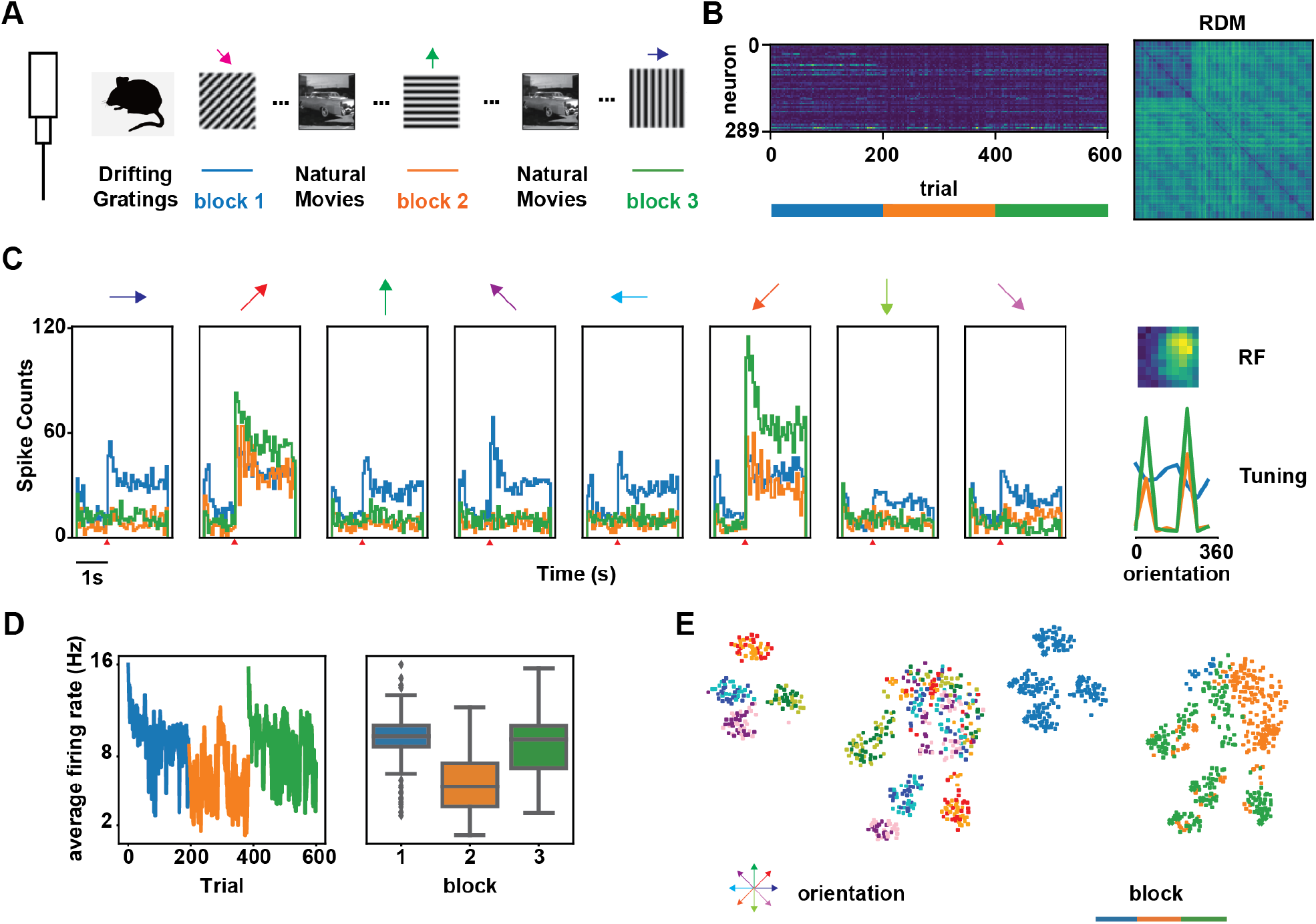
Neural representational drift causes embedding shift across blocks of drifting grating stimulus. **A-E:** An example session of Neuropixel electrophysiological recording (750749662). **A**. Schematic illustration of Neuropixel recording. **B**. Population response matrix (middle, *n* = 289) and representational dissimilarity matrix (RDM) as pairwise distance between population vectors. We first ordered the trials by block. And within each block, we ordered the trials according to the stimulus direction. **C**. Block modulation of the response of an example neuron ( # 951867364). *left*, peri-stimulus time histogram (PSTH) per block and direction. *upper right*, receptive field of the neuron, estimated by the neural response to Gabors at different positions. *lower right*, direction tuning curves per block of the neuron. For this specific neuron, the direction-tuning curve was sharpened in the second and third blocks. **D**. Overall neural response of the visual cortex to each drifting grating trial. Activity was defined as the mean spike rate of all units recorded in the visual cortex within 200 ms after stimulus onset. The general neural activity of each block was summarized as the average activity of all the trials in the block. **E**. t-SNE embedding based on RDM. *upper*, labeled by the directions of the drifting grating stimuli. *lower*, labeled by block.

To quantify the representational drift between blocks, we built a simple linear classifier to classify the block to which the trial belonged to. In Neuropixel recordings, we found that for 78.1% of all sessions (N=32), the cross-validated accuracy of a trained block classifier was significantly higher than the chance level (p-value ≤ 0.005), and the average accuracy was about 54%. Furthermore, the average inter-block distance between trials was significantly greater than the average distance within the same block (Figure S1C).

Although drift appears to be substantial in the low-dimensional space, in the high-dimensional neural state space, the decoder built from one block generalized relatively well on the other blocks (Figure S3A-C). A further investigation into the weights of the linear classifier showed that the decision boundary of the block classifier is close to perpendicular to that of the orientation classifier (Figure S3D). Thus, representational drift across blocks is orthogonal to the encoding plane of the orientations, similar to what was previously reported in other data (Aitken et al., 2022).

Further analyses comparing different probes and calcium data, in which similar block effects were found, suggest that these drift effects did not reflect the physical movement of the electrode through the brain tissue (Figure S4). To investigate the possibility that drift reflects behavioral dynamics, we performed various control analyses either by regressing out behavioral covariates; computing representatial drift for distinct behavioral states; or via Tensor Component Analysis (TCA). These analyses, together with the results on the time courses of representational drift presented below, suggest that the observed representational drift was not due to behavior (Fig. S5, Fig. S7).

Finally, we observed that loadings onto drift TCA components did not differ across visual areas and were uncorrelated with laminar depth, indicating that representational drift was found across areas and layers (Fig. S8A–B).

### Representational drift shows same-class-acceleration

Next, we analyzed the speed of representational drift both across the three blocks and within (i.e inter-block) the three blocks. To quantify the speed of representational drift, we calculated the correlation between population responses as a function of the time interval *τ* between different trials corresponding to the same stimulus. This defined a Population Vector Correlation (PVC) between two population vectors as a function of 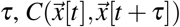. Because of the structure of the session, the time intervals can essentially be divided into three regions: The first region contains the shorter time intervals corresponding to the drift within a single block; the second region contains the time intervals between blocks 1-2 and 2-3 (i.e. the inter-block intervals); the third region contains the time interval between blocks 1 and 3.

To understand what kind of drift patterns may emerge from varying the drift speed within and between the blocks, we first performed simulations. We simulated a population of neurons with orientation tuning curves. Each neurons firing rate was modeled as a stochastic random walk, where a random innovation was added to its tuning curve for each trial (Figure S5A). Using this basic model of representation drift, we simulated four different drift patterns: (1) continuous drift with a constant speed; (2) only within-block drift; (3) only inter-block drift; and (4) drift with deceleration during the inter-block intervals (See Methods). We then computed the change in the PVC as a function of *τ*. Figure 3A showed that, with different speed patterns, PVC curve exhibited different slope patterns. In the “continuous” condition (1), the slope remained steady, producing a linear pattern. In the “only within” condition (2), the PVC curve declined more rapidly within each block and recovered during the inter-block gap, resulting in an “M-shaped” slope pattern. In the “only inter-block” condition (3), the curve remained flat within blocks and decreased only during the inter-block gap, creating a “W-shaped” slope pattern. In the “fast within, slow inter” condition (4), the slope pattern exhibited a similar “M-shaped” profile as observed in the “only within” condition. However, due to the slower drift during the inter-block intervals, the PVC curve did not recover as strongly, resulting in slightly negative slopes between blocks, and a flatter “M” shape.

**Figure 3.**
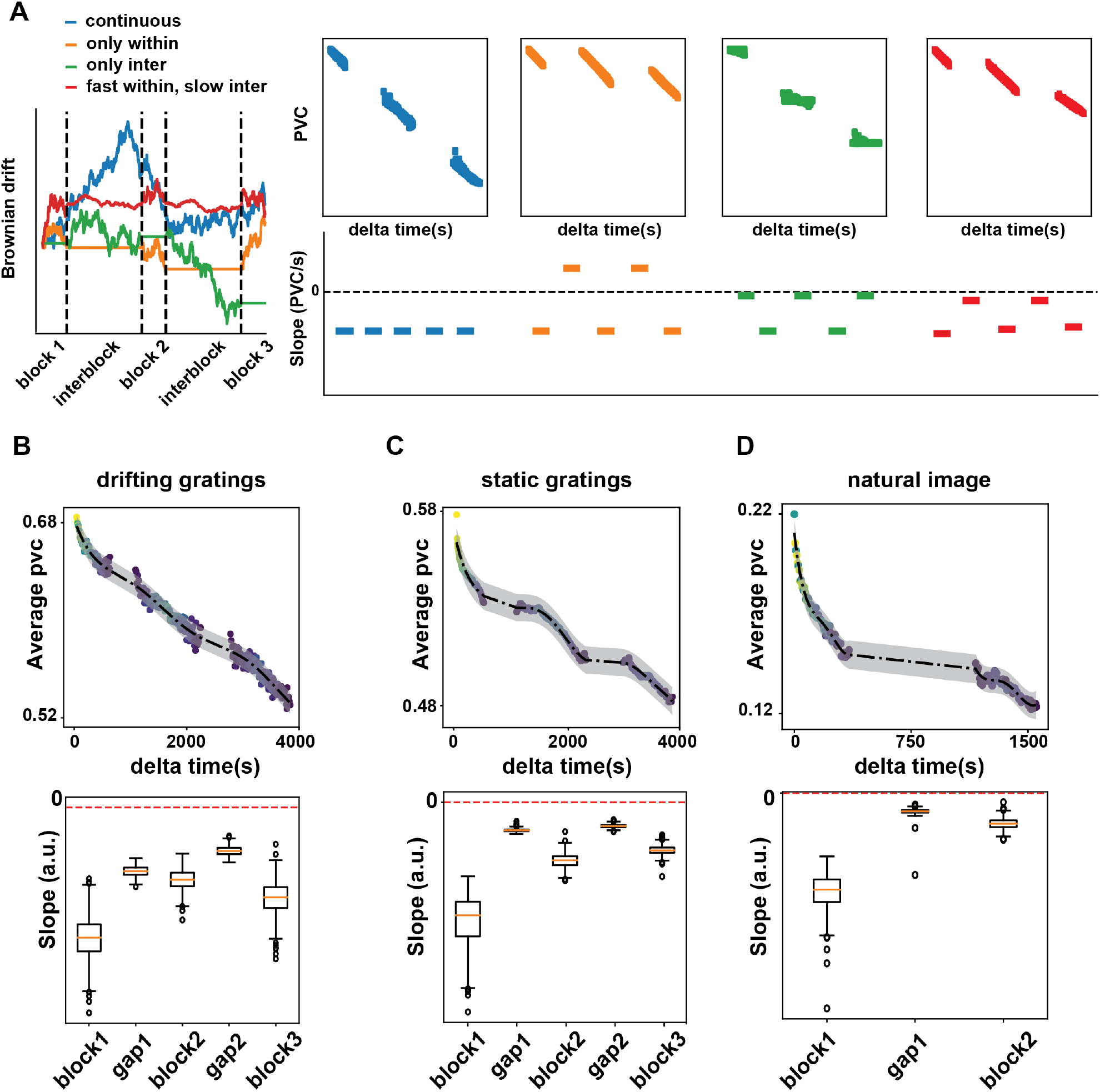
Representational drift is independent of stimulus type and is continuous regarding population vector correlation. **A**. Simulation results of the PVC curve. Representational drift was modeled as independent Brownian motion of neuron population in three different ways. Blue: Continuous Brownian motion with constant velocity. Orange: Brownian motions occur only during stimulus presentation. Green: Brownian motion occurs only between stimulus blocks. Red: Brownian motion is faster within blocks and slower in inter-block duration. **B**. Average population vector correlation as a function of Δ*T* across all animals (*n* = 32) for drifting gratings. **C**. Average population vector correlation as a function of the time interval between two static grating trials with the same orientation. **D**. Average population vector correlation as a function Δ*T* of trials that respond to the same natural image. All population vectors are normalized according to the running speed to remove the influence of behavior (See Method ).

We computed the PVC to investigate the structure of drift in the neural recordings. Visualization of the average PVC as a function of *τ* across sessions suggests that the correlation between population responses decreased monotonically as a function of the time-interval *τ* (Figure 3B). That is, population vectors continue to decorrelate as time elapses within a session. The slopes for the five intervals were significantly negative and exhibited a characteristic “M-shaped” pattern (Figure 3B). This result indicates that there is faster representational drift within the blocks, and that the drift slows down during the inter-block intervals. Consistent with the analyses presented above, the drift remained significant during the inter-block intervals.

A similar patterns were observed for static gratings (see Methods), in which the blocks were interspersed with mainly natural scenes (block 1-2) and movies / scenes (block 2-3) (Figure S5B-C). A similar pattern was also observed for natural scenes, which were interspersed by a block of stating gratings and natural movies. We thus conclude drift in neural representations of stimuli of class “A” (e.g. drifting gratings) accelerates when presenting stimuli of that class (e.g. drifting gratings), i.e. within the blocks; while drift decelerates when presenting stimuli of other classes.

### Representational drift is accompanied by noise correlation reshaping

If representational drift is partially driven by experience-dependent changes in synaptic weights, then we would expect the synaptic strength within the neuron ensemble to reorganize across blocks, leading to connectivity reshaping and representational drift. Therefore, we next investigated the functional connectivity of the neural population. We found that indeed the noise correlation of the neuron ensembles reshaped across blocks (Figure 4A). We measured the similarity between the noise correlation matrices of any two blocks as the correlation coefficient between their off-diagonal elements. This correlation, termed pattern correlation, reflects how much the noise correlation pattern reshapes (Hofer et al., 2011). A larger pattern correlation means that the noise correlation pattern between units shows less reconfiguration. We found that the noise correlation pattern changed most between the first and the last blocks and such change was mainly due to the reshaping between the first two blocks (Figure 4B). Moreover, the strength of the noise correlation reconfiguration was correlated with the strength of representational drift (Figure 4C), suggesting that the representational drift was accompanied by synaptic structure reconfiguration of the neuronal ensemble.

**Figure 4.**
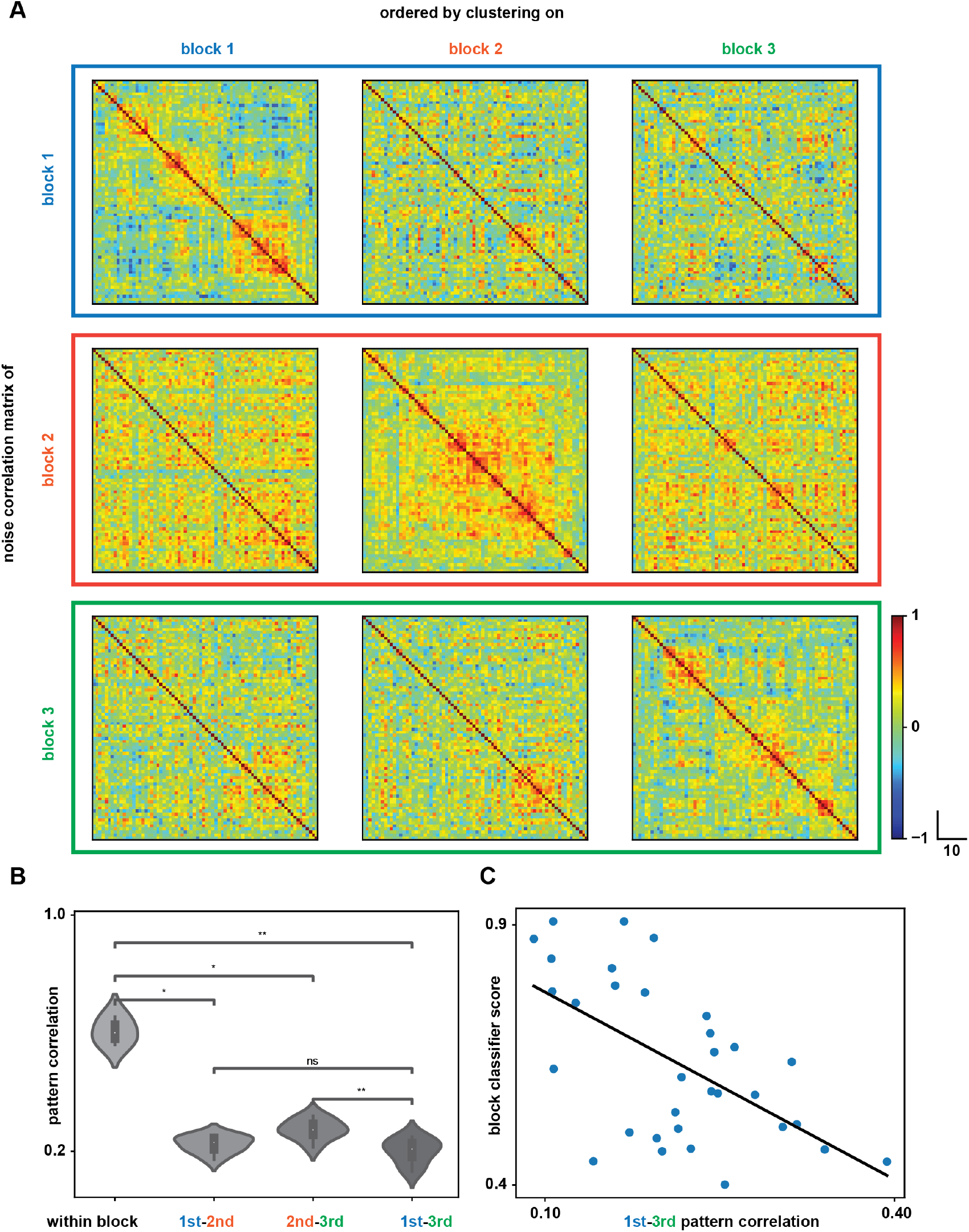
Representational drift and noise correlation reshaping. **A**. Pairwise noise correlation matrices of aneural ensemble response to a given visual stimulus (orientation = 315^*°*^) across three blocks. On each row is the noise correlation matrix of each block. The ordering of neurons is the same for each column based on hierarchical clustering. **B**. Pattern correlation between any two blocks for all stimulus orientation and all animals. **C**. The average pattern correlations between the noise correlation matrices of block 1 and block 3 are negatively correlated with the cross validated score of SVM classifier trained to classify block labels.

## DISCUSSION

### Summary

Representational drift refers to incremental changes in neural responses to the same stimulus across repetitions. Representational drift within a session likely reflects a combination of numerous mechanisms including changes in behavioral state, adaptation, fluctuations in excitability and synaptic plasticity (Micou and O’Leary, 2023; Sadeh and Clopath, 2022; Rule and O’Leary, 2022; Attardo et al., 2015; Rule et al., 2020; Delamare et al., 2024). Here, we analyze multi-areal electrophysiological recordings to identify representational drift within a session in visual cortex. Because the sessions were organized in different blocks, the low-dimensional embedding showed relatively large shifts in representations across blocks due to the passage of time between blocks. We show that this representational drift is not explained by changes in behavioral state and has a specific temporal structure that reflects the structure of sensory experience rather than behavior. Specifically, we found that the speed of representational drift for a specific stimulus category was not constant across trials, but showed acceleration during the blocks in which those stimuli are presented, and deceleration during blocks in which other stimuli were presented. Using theoretical analysis and simulations, we show that this behavior is predicted from experience-dependent plasticity: learning induced by a stimulus of category *c* should produce larger changes in neural representations for test probes that are more similar in population space. This same-class acceleration principle provides a general strategy to dissociate different sources of drift experimentally.

### Potential influence of behavior

It is well established that the behavioral state of an animal acts as a brain-wide background signal contributes substantially to changes in neural representations (Niell and Stryker, 2010; Stringer et al., 2019; McGinley et al., 2015; Vinck et al., 2015). Thus, representational drift may be partially explained by changes in behavioral state within a session (Sadeh and Clopath, 2022). Our analyses suggest that representational drift can only be partially explained by changes in behavioral state: First, we performed several control analysis by separating running and sitting periods, as well as analyzing representational drift based on regression residuals. Second, Tensor Component Analysis TCA) showed that representational drift was a separate factor from the behavioral factors, and showed a clear correlation with the block structure of the session, in contrast to the behavioral factors. Third, we showed that representational drift showed a systematic temporal structure that cannot be explained by changes in behavioral state alone: Drift velocity was highest during the stimulus blocks in which the stimulus was presented, and had a lower velocity in between blocks. Notably, we found that differences in behavior as a function of time did not mimick this pattern.

### Representational drift and changes in synaptic connectivity

Representational drift within a session may be caused by continuous fast learning (Qin et al., 2023; Hennig et al., 2021; Schoonover et al., 2021; Bauer et al., 2023). Recent work suggests such learning can take place in a self-supervised manner and may not differ substantially between passive exposure and task learning (Zhong et al., 2025). Such self-supervised learning may take the form e.g. of predictive coding (Aizenbud et al., 2025; Uran et al., 2022), but could also be consistent with a discriminative neural network trained via unsupervised loss functions (Nayebi et al., 2023). The input-dependent hypothesis on representational drift is consistent with the findings reported here: First, it is consistent with the structure that we observed, namely of accelerated drift in stimulus responses during the blocks in which that stimulus is presented, with decelerated drift in between blocks. Second, it is consistent with the changes in pairwise neural correlations across blocks.

Our central empirical observation is that representational drift within a session is input-dependent: the drift speed for a stimulus class accelerates during blocks in which that class is presented and decelerates otherwise. The theory developed here predicts exactly this pattern. A single SGD step driven by an exemplar of class *c* produces a representational update for a probe *x* whose magnitude is proportional to a kernel similarity between *x* and *x*_*c*_, measured in the current representational space. Thus, repeated exposure selectively increases drift for stimuli that already project strongly onto the class-c, i.e., same-class-acceleration, while other classes are affected less. This provides a mechanistic account for the block-wise acceleration/deceleration we observed.

Interestingly, we found that drift took place predominantly in a subspace orthogonal to the coding subspace, leading to little degradation for classifiers across blocks, which matches previous conclusions (Aitken et al., 2022). Aitken et al. (2022) explain this from noise in SGD updates in the training of a neural network. The orthogonality can be expected to be larger late in the training of a neural network; gradients along task-relevant, high-curvature directions will be smaller, and gradients tend to accumulate in flatter directions. Furthermore, one expects orthogonality more in earlier layers of a neural network. Same-class acceleration of representational drift however should persist later in training, and we show that it becomes stronger as feature similarity within the same class gets stronger. Thus, the findings of same-class-acceleration and subspace orthogonality may suggest that the visual cortex shows characteristics of a network in the “late training” phase. This would predict that same-class-acceleration is substantially weaker in early development, while the representational drift subspace shows less orthogonality w.r.t. stimulus space.

Recent longitudinal imaging shows that sensory experience biases drift direction in V1 across weeks towards the repeated stimulus, consistent with experience-dependent plasticity and the same-class-acceleration principle proposed here (Bauer et al., 2023). At the same time, recent work has shown stimulus-specific plasticity of gamma synchronization and firing-rate repetition effects in macaque V1/V2; critically, these changes accumulate across time but transfer only weakly across different stimuli (Peter et al., 2021; Brunet et al., 2014; Psarou et al., 2025). That pattern can be explained by the same-class acceleration principle: repetition of the same stimulus changes synaptic weights which lead to a change in network oscillations specifically for that stimulus; these changes in synaptic weights however weakly affect gamma-band responses for other stimuli. Our findings also agree with a large body of behavioral work on perceptual learning that shows limited transfer beyond the trained features/tasks (e.g., orientation, retinal location) (Krakauer et al., 2006; Lange et al., 2018; Fahle, 2005).

It stands to reason that continuous representational drift may in addition be caused by independent random synaptic turnover without being induced by learning. In the mouse cortex, more than 60% of synapses are replaced within three weeks (Loewenstein et al., 2015) and much of the synaptic reorganization takes place spontaneously (Mongillo et al., 2017). The hippocampus completely reorganizes within one month (Attardo et al., 2015) and more than half of the fluctuations are attributed to spontaneous processes rather than learning (Dvorkin and Ziv, 2016). These observations led to the proposal that a random walk in the weight space can lead to spontaneous reshaping of the connectivity, and thus to the drift in the neural representational space.

In sum, representational drift within a session likely reflects a number of mechanisms, including fluctuations in behavior and excitability, input-dependent plasticity and random synaptic turnover. We have shown here that some features of representational drift are consistent with a role for input-dependent plasticity, as the time course of drift varies with sensory experience and shows same-class-acceleration. These are theoretically predicted features of experience-dependent weight updates in a neural network trained by gradient descent.

## METHOD

### Electrophysiological recording

Data were downloaded in NWB format from the Allen Neuropixel Brain Observatory. There are in total 32 adult mice subjects in brain observatory 1.1 paradigm with an average age of 112 days. 27 of these mide are male and 5 are female. 16 of the mice are wild-type mice, and the rest come from transgenic lines (*N* = 6, Sst-IRES-Cre/wt; *N* = 5, Pvalb-IRES-Cre/wt; *N* = 5, Vip-IRES-Cre/wt). Rodents were head fixed, and presented with different types of visual stimulus in a certain order. In total, six high-throughput Neuropixel probes were inserted into mouse brain and recorded simultaneously from both visual cortical areas and subcortical areas including hippocampus and thalamus. For visual neurons, we include only the units with signal-to-noise ratio larger than 0.5. Moreover, in order to rule out the influence of possible physical probe drift, we further discarded neurons that exhibited a sudden silencing during the session (Cowley et al., 2020), which is defined as the lack of spiking activities for consecutive 20 trials regardless of the presented stimuli. To further screen for sensory sensitive visual neurons, we selected units with a strong overall orientation tuning and a clear receptive field (See Supplementary S1). We measure the orientation selectivity of a single neuron by carrying out one-way ANOVA test on its response to drifting gratings of eight directions. We selected units with strong tuning preference (p-value *<* 1e-3). We estimated the receptive field of a single neuron by its average response to Gabor stimuli. Gabors were shown on 81 different positions in a random order. Each Gabor stimulus is repeated for 45 times. Using ANOVA, we measured the variability that is explained by the stimulus position, and we only included neurons with clear position clustering (*µ >* 0.05).

After unit selection, we included only sessions with more than 30 neurons to have a robust estimate of the manifold. Using AllenSDK python API (version==2.13.1), we counted the number of spikes of selected *N* neurons within a window of 200 ms after the stimulus onset as the population response. The short time window was chosen to study the fast temporal dynamics of visual response. This yields population vector for each trial *X*_*i*_ ∈ ℝ^*N*^, ∀*i* = 1, …, 600. To calculate each entry in the representational dissimilarity matrix (RDM), we calculated the Euclidean distance between population vectors for any pair of trials 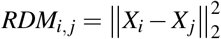.

### 2p calcium imaging

Data were downloaded in NWB format from the Allen Institute Brain Observatory. 540 transgenic mice (N = 76 Cux2-CreERT2, N = 41 Emx1-IRES-Cre, N = 9 Fezf2-CreER, N = 39 Nr5a1-Cre, N = 18 Ntsr1-Cre GN220, N = 21 Pvalb-IRES-Cre, N = 38 Rbp4-Cre KL100, N = 40 Rorb-IRES2-Cre, N = 9 Scnn1a-Tg3-Cre, N = 100 Slc17a7-IRES2-Cre, N = 64 Sst-IRES-Cre, N = 9 Tlx3-Cre PL56, N = 76 Vip-IRES-Cre) expressing GCaMP6f were imaged through cranial windows with different laminar depth and six visual cortical regions.

Using Suite2p package (Pachitariu et al., 2017), we computed deconvolved Δ*F/F* values in the two seconds after the stimulus onset. Compared with the analysis of electrophisiological data, a longer window was used considering the relatively slower temporal dynamics of calcium imaging.

### t-SNE visualization of low-dimensional neural manifold

To visualize the manifold of neural representations of visual stimuli, we used t-distributed Stochastic Neighbor Embedding (t-SNE, Van der Maaten and Hinton (2008)) as a dimensionality reduction method. Representational dissimilarity matrix (RDM) was computed as the pairwise Euclidean distance between population vectors of either electrophysiological or calcium imaging datasets. Based on the computed RDM, we approximated the proximity of data points in the low-dimensional 2D embedding with t-SNE using a perplexity value of 10.

### Tuning curve modulation analysis

To measure the block-specific tuning curves of visual cortical neurons to drifting gratings, we computed the average spike counts within 200 milliseconds after the stimulus onset of each neuron in three blocks. To model both orientation and direction tuning, we fitted the bimodal curve by a sum of two von Mises functions with different preferred orientations 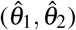, amplitudes (*a*_1_, *a*_2_), scales (*b*_1_, *b*_2_) and bandwidths (*κ*_1_, *κ*_2_) (Graf et al., 2011; Arandia-Romero et al., 2016).

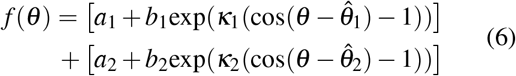

And to quantify the change of orientation tuning curves between blocks, we performed linear regression on von Mises fits of two different blocks, *f*_*i*_(*θ*) = *α f* _*j*_(*θ*) + *β* . The slope of the fitted regression model *α* was termed as Multiplicative Factor (MF) and the intercept *β* was termed as Additive Factor (AF). As a control, we measured tuning curve of high response vs tuning curve of low response this is to illustrate that the representational drift is not simply due to global scaling To eliminate the correlation between mf and af: estimate af from odd trials, estimate mf from even trials

### Quantification of representational drift by block decoder

To quantify the representational drift as a shift of manifold, we built a naive linear classifier to classify the block label. Here, we used Support Vector Machine (SVM) implemented by [sklearn](https://scikit-learn.org/stable/modules/classes.html#module-sklearn.svm). As expected, after shuffling the block label, the performance of the classifier went down to a chance level.

### Quantification of sensory encoding stability

To examine how orientation decoding performance was influenced by representational drift, we built SVM decoders to distinguish trials of different orientations. We used 5-fold cross-validation to measure the generalization.

### Regression from behavior to neural response

To control for the influence of animal behavior, we regressed each neuron’s response **x**_*i*_ against its arousal signal measured as the pupil diameter *P* and locomotion signal measured as the running speed *R*. Combining both behavioral signals as dataset *B*, the coefficients of the ridge regression model **w**_*i*_ was estimated by minimizing the sum-of-squared error with a regularization term.

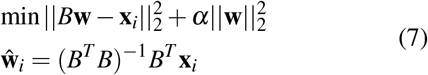

By fitting the regression model for each neuron *i* and keeping only the residual of the regression model *ε*_*i*_, we isolated the behavior factor from the variability of interest.

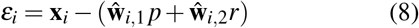

### Disentangle representational drift by tensor component analysis

We used Tensor Component Analysis (TCA, Williams et al. (2018)) to find the component that captures the representational drift. Since the number of repetitions for each stimulus condition is the same (*K*) and the ordering of stimulus was randomized, therefore, we can reorganize the neural activities as a fourth-order tensor *X* ∈ℝ^*N×T×S×K*^, where *N* is the number of neurons, *T* is the number of time points, *S* is the number of conditions and *K* is the number of repetitions for each condition. The unsupervised dimensionality reduction algorithm TCA decomposes the dataset *X* into a sum of *J* rank-one tensors. Each rank-one tensor is the outer product of four tensor components.

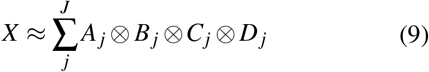

The trial component *D*_*j*_ ∈ℝ^*K*^ summarizes the neural trial-by-trial variability and representational drift. Factorization of 4D neural tensor was implemented by using the non-negative PARAFAC algorithm from tensorly https://tensorly.github.io/. We chose the number of components of TCA to be the smallest dimension of the tensor (*J* = min {*N, T, S, K*} ).

To identify the trial components that are drift-related, we computed the Pearson Correlation Coefficient between the trial components and the pupil diameter, corr(*D*_*j*_, *P*). To exclude the components that encode the visual signal, we calculated the Orientation Selectivity Index (OSI, Swindale (1998)). Therefore, we include only the factors that have small OSI (OSI(*C*_*j*_) *<* 0.7) and low correlation with behavioral signal (corr(*D*_*j*_, *P*) *<* 0.5). To compute the dissimilarity matrix of the drift components, we randomly initialized the TCA model ten times, gathered all the identified drift components, and calculated the distance between repetitions with each drift component as a feature.

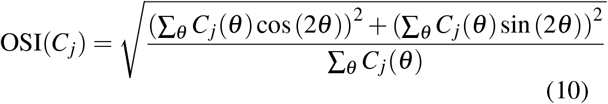

### Cluster trials with hidden markov model

We used a Hidden Markov Model (HMM) to identify highly aroused period and also up state during spontaneous activity. HMM is a generative model that assumes the observed sequence to be generated from a sequence of discrete hidden states. By fitting a HMM, it learns the transition probability between states and the emission probability between hidden states and observed variables. Here, we choose Gaussian as the emission process. By fitting a two-state model to the pupil diameter data, we can cluster the trials into high/low arousal states. Similarly, by fitting a two-state model to the animal running speed, we can cluster the trials into running and quiescent states. Then, fitting another three-state HMM to the population spiking activities in the quiescent period, we can identify the up state during the spontaneous activities.

### Noise correlation analysis

To measure the noise correlation between neurons, we computed the correlation between responses of any pair of neuronal units to the same stimulus. To find groups of neurons that co-fluctuate, we adopted an agglomerative hierarchical clustering algorithm. The clustering ordering of the neurons was calculated using UPGMA (Unweighted Pair Group Method with Arithmetic mean). To compute the pattern correlation between any two noise correlation matrices, we measured the correlation between their off-diagonal elements. Average pattern correlation was calculated across stimuli of all directions and across all animals. As a control, we calculated the within-block pattern correlation by randomly splitting each block into two halves and computing the noise correlation matrices respectively for each half.

### Quantification of representational drift by block modulation index

To quantify the latent variables that explain the representational drift between blocks, we computed the block modulation index (BMI) as the largest difference between mean values of any pair of blocks, normalized by their sum. The latent factor was made non-negative before computation. Suppose *Z* is a non-negative factor, 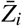 is the mean value in *i*^th^ block, and 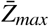 and 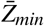 are the largest and smallest mean value among all blocks. Then, BMI is calculated as the following,

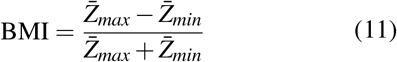

A factor with no block modulation would have BMI=0, and the strongest modulation would have BMI=1.

### Simulation of representational drift by Brownian Motion

We simulated representational drift using independent Brownian motion. First, we generated a population of 500 neurons with tuning curves evenly distributed across the orientation space. We presented stimuli of eight different orientations to this neuronal population across three blocks. Each trial lasted 2 seconds and was repeated 20 times within each block. Between adjacent blocks, there’s a 1000-second interval. Each neuron independently accumulated Brownian noise into its firing rate at each time point. We examined four different conditions for the noise accumulation rate. In the first condition, termed “continuous,” the noise accumulation rate was constant across the entire simulation. In the second condition, “only within,” Brownian motion stopped during the inter-block intervals, occurring only within blocks. In the third condition, “only inter,” Brownian motion occurred exclusively during the inter-block intervals. In the fourth condition, “fast within slow inter”, Brownian motion is relatively fast within the blocks and slow during the inter-block intervals.

### Theoretical analysis

Here, we provide systematic analyses of the change in representations across time due to training a neural network via stochastic gradient descent. We shall do this for the case of logistic regression, the readout layer in a multi-layer neural network, and any layer of a neural network.

#### Case of logistic regression

Let the input feature vector be *x* ∈ ℝ^*d*^ . The classifier has weight parameters *W* ∈ℝ^*K×d*^ and biases *b* ∈ ℝ^*K*^, logits *z*(*x*) = *Wx* + *b* ∈ ℝ^*K*^, probabilities *p*(*x*) = softmax(*z*), and per-example loss ℒ (*x*) = − log *p*_*c*_(*x*) when the label is *c*. Consider one stochastic-gradient step of size *η >* 0 for one input from class c, *x*_*c*_.

We measure the change in a test probe’s response before vs. after the weight update. Let *t* index the parameter state before the step and *t*+1 the parameter state after the single SGD step for an input *x* of class *c*. For any probe (test) input *x*^′^, we define

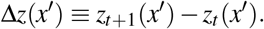

We will compare ‖ Δ𝓏 (*x*^′^)‖_2_ for probes with *x*′ ∈ *c* versus *x*^′^ ∉ *c*.

A single SGD step of size *η >* 0 for input of class *c* causes a weight update

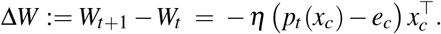

Here, *e*_*c*_ denotes the class-*c* one–hot target vector in ℝ ^*K*^ . For any probe *x*^*′*^,

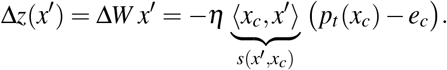

Hence, the change vector is a fixed direction set by the training example’s *p*_*t*_(*x*_*c*_) − *e*_*c*_, scaled by the scalar similarity *s*(*x*′, *x*_*c*_).

Hence

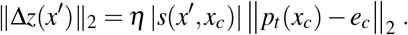

The factor ‖*p*_*t*_(*x*_*c*_) *− e*_*c*_‖_2_ is fixed by the single training example; therefore, for any two probes 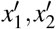,

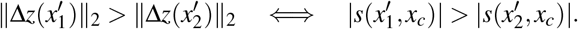

In particular, if probes from class *c* are, in expectation, more similar to *x*_*c*_ than probes not in *c*, i.e.

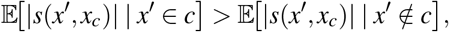

then

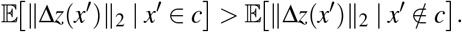

Finally, we can consider the special case of a test probe which is the training sample itself. If we consider unit-norm feature vectors, then the maximal change in representation will occur when *x*^′^ = *x*_*c*_.

### Case of multi-layer neural networks

We use pre-activations 𝓏 ^(𝓁)^( ·) and activations *a*^(𝓁)^( ·) with a nonlinearity *h*.

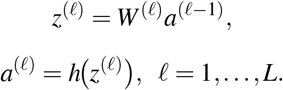

Readout and probabilities are defined as

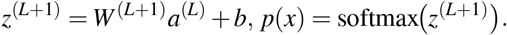

Let the loss be ℒ (*x, y*) (e.g. cross-entropy). We analyze one SGD step of size *η >* 0 on (*x*_*c*_), an input from class *c*, and its effect on the layer-𝓁 activation *a*^(𝓁)^(*x*^′^) of any probe *x*^′^ .

Only parameters up to layer 𝓁 can directly change *a*^(𝓁)^: *θ*_*≤* 𝓁_ ≡ {*W*^(1)^, …,*W*} ^(𝓁)^ .

We define the pre-activation Jacobian

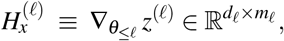

and the backprop delta error signal

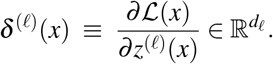

Note that for Cross-entropy loss: *δ*^(*L*+1)^(*x*) = *p*(*x*) *− e*_*y*_, and *δ*^(𝓁)^ (*x*) = (*W*^(𝓁+1) *T*^*δ*^(*f*+1)^ (*x*)) ⨀ (*h*′ (𝓏^(𝓁)^ (*x*)) for 𝓁≤ *L*. Define the layer-𝓁 pre-activation kernel and the normalized unit class-*c* gradient direction:

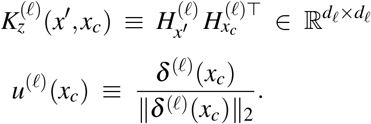

Note that 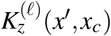, when defined at the level of logits, is the Neural Tangent Kernel (NTK) that is used to describe the evolution of neural networks in the infinite-wide limit. The update in weights due to *x*_*c*_ equals

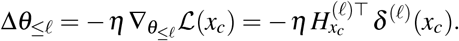

For the probe *x*^′^, we can write the layer-𝓁 preactivation as a function of the parameters up to that layer:

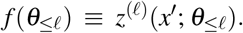

For a small step Δ*θ*_*≤*𝓁_ around *θ*_*≤*𝓁_,

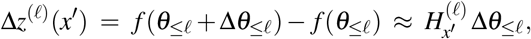

where 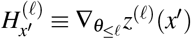.

For any test probe *x*^*′*^ we thus have the first-order approximation

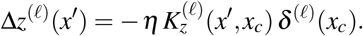

This yields the change in neural representation

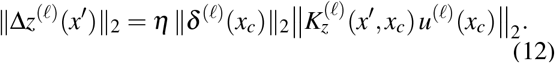

Define the (probe-dependent) similarity score

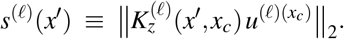

which determines the change in neural representation depending on the class-identity of the probe *x*^′^ . Then (12) implies that, for fixed (*x*_*c*_, *c*), comparing ‖ Δ𝓏^(𝓁)^(*x*^*′*^)‖_2_ across probes is equivalent to comparing *s*^(𝓁)^(*x*^*′*^). In particular, if

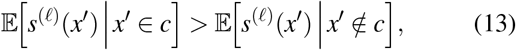

then

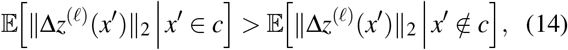

and, up to local *h*′ factors, this statement holds for ‖ Δ*a*^( *𝓁*)^(*x*^′^)‖_2_.

### Logits in readout layer

We first consider the behavior of the **logits in the readout layer** in a trained neural network, where we consider only an update of the weights in the last layer. Let 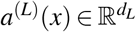 be the last hidden activation of a ReLU network and

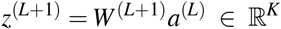

be the logits (note we hold *b* fixed). With cross-entropy loss, for an input-label pair (*x*_*c*_, *c*) we have

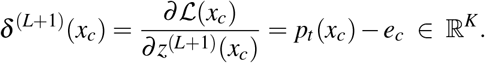

The gradient w.r.t. the readout weights is the product

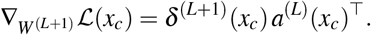

A single SGD step of size *η >* 0 gives

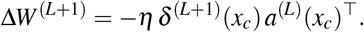

For any probe *x*^′^, the change in its logits is therefore

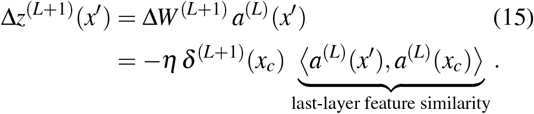

Taking norms,

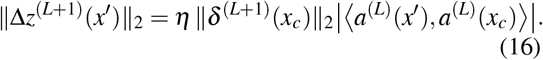

Equation (16) shows that, for fixed (*x*_*c*_, *c*), the probe dependence is via the scalar similarity *s*(*x*^*′*^) ≡ ⟨*a*^(*L*)^(*x*^′^), *a*^(*L*)^(*x*_*c*_)⟩ of last-layer features. In trained ReLU networks, within-class inputs typically share gates ReLU gates and cause activations that cluster in feature space, which increases the inner product ⟨*a*^(*L*)^(*x*^′^), *a*^(*L*)^(*x*_*c*_) ⟩ for *x*′ ∈ *c*. Under these assumptions,

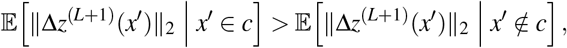

establishing same-class-accelerated drift in the readout layer. Again, among probes with fixed ‖*a*^(*L*)^(*x*)‖_2_ (e.g., under LayerNorm or *L*^2^-normalization), the update magnitude is maximized in the special case that the test probe equals the training probe, i.e. *x*^*′*^ = *x*_*c*_, where

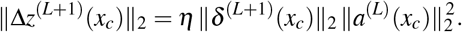

### Relation to Neural Tangent Kernel

Note there is a relation to the Neural Tangent Kernel. Recall the definition of the pre-activation Jacobian and backprop deltas. For logits, set 𝓁 = *L*+1 and let *θ*_*≤L*+1_ ≡ *θ* be all trainable parameters. Then

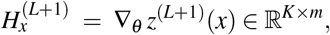

and

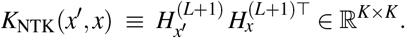

is the well-known Neural Tangent Kernel (NTK) which analytically describes the evolution of network outputs in the infinite-width limit. In the NTK (infinite-width) limit, the network is globally linearized around initialization: the Jacobian 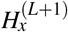 (hence *K* _NTK_ ) is constant during training, and a single SGD step on (*x*_*c*_, *c*) therefore produces the exact update

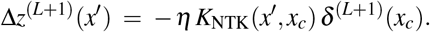

### Neural network representations in any layer

Next, we consider the case of representations in any layer of the neural network, starting from eq. 12, with the relevant quantity to consider 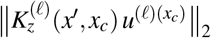. Recall that *θ*_*≤* 𝓁_ = {*W*^(1)^, …,*W*^(𝓁)^}. We can write the Jacobian as a concatenation over layers:

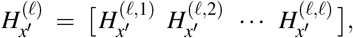

where each block is the Jacobian w.r.t. *W*^( *j*)^:

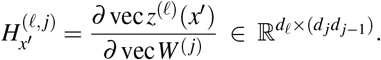

We define the inter-layer Jacobian

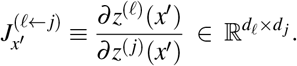

Hence

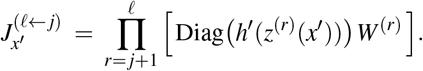

We thus obtain

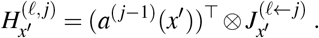

The second component of 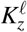 can be written as

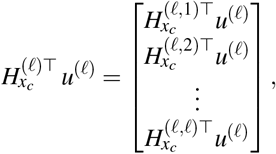

with the *j*-th block is 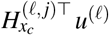. Note that

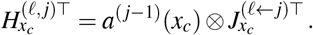

Define 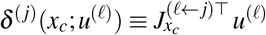. Note that substituting *u*^(𝓁)^ yields the exact identity

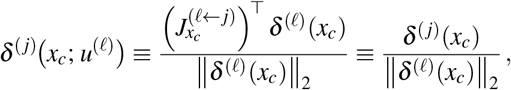

i.e. the standard backprop signal for layer *j* scaled by the magnitude of the backprop signal for layer 𝓁.

We can now write

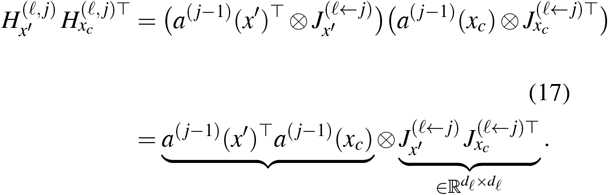

Hence we can write

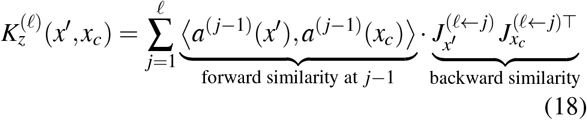

We can thus see that 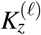 depends on the similarity of activations up to layer 𝓁. However, 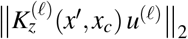 depends also on the term 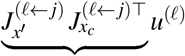, which cannot be expressed as a similarity measure. However, we can express a lower bound explicitly in terms of forward and backward similarities.

Define

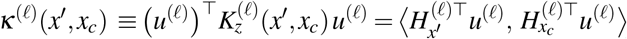

From this we obtain a Cauchy-Schwarz bound

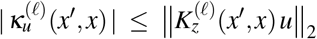

We can see that

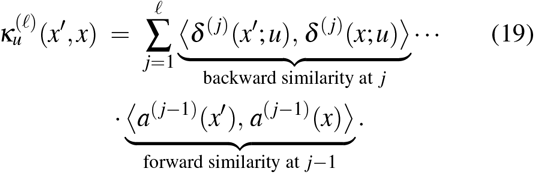

Let *x*_*c*_ be a training example of class *c*, and let *u*^(𝓁)^ ≡ ‖ *δ*^(𝓁)^(*x*_*c*_)*/ δ*^(𝓁)^(*x*_*c*_)‖_2_ be the unit class-*c* gradient direction at layer *f*. Training in ReLU nets promotes (i) forward clustering *a*^( *j−*1)^(*x*) ≈ *a*^( *j−*1)^(*x*_*c*_) within class (helped by shared gates) and (ii) backward alignment of the deltas *δ*^( *j*)^(*x*^*′*^ ; *u*^(𝓁)^) with *δ*^( *j*)^(*x*_*c*_; *u*^(𝓁)^) along class-discriminative directions. Again, if the test and train probe are identical, then one expects 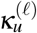 to be maximal if we assume normalized network activations.

As depth inside the neural network increases, the forward features *a*^( *j−*1)^( · ) are progressively denoised and clustered. ReLU gates become more similar within class, so ⟨*a*^(*j−*1)^(*x*^*′*^), *a*^(*j−*1)^(*x*_*c*_) ⟩ grows for *x*^′^ ∈*c*. Backward signals *δ*^( *j*)^(*x*_*c*_; *u*^(𝓁)^) and *δ*^( *j*)^(*x*^*′*^ ; *u*^(𝓁)^) share similar gates and weights for same-class inputs, which boosts ⟨*δ*^( *j*)^(*x*^*′*^ ; *u*^(𝓁)^), *δ*^( *j*)^(*x*_*c*_; *u*^(*𝓁*)^) ⟩ . Consequently, the sum in (5) should exhibit stronger same-class changes at deeper layers on average.

Thus, for pre-activations at any layer, the directional kernel 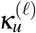 decomposes into a sum of (gated) forward *×* backward similarities. Cauchy–Schwarz inequality turns this decomposition into a lower bound on the actual response magnitude 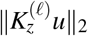. Since training increases both forward feature similarity and backward alignment within class-especially in deeper layers—this yields a theoretical explanation for same-class acceleration that becomes more pronounced with depth and training.

### Neural Networks simulations

#### Logistic Regression on MNIST

To empirically validate our theoretical framework, we began with a logistic regression model trained on the MNIST digit classification task. The input pixel values were first normalized and reduced to 20 principal components using PCA. The model was then trained sequentially, one digit class at a time. After each class-specific update, we evaluated the model on the entire test set and computed the mean output logits for each class **z**_*t*_. Changes in these mean representations were quantified using the L2 distance between successive training steps: Δ**z**_*t*_ = ‖**z**_*t*+1_ *−* **z**_*t*_ ‖_2_.

#### Covolutional Neural Network on CIFAR10

Next, we trained deep convolutional neural networks (CNNs) on an image classification task. We used Cifar-10 dataset that contains 60000 32 *×* 32 color images in 10 object classes (50000 training images and 10000 test images). We created a deep CNN formed of 3 convolutional layers, each followed by a max-pooling down-sampling layer. The representations of the last convolutional layer are then vectorized and followed by a 64-dimensional fully-connected penultimate layer before the classification layer. The representations of the fully-connected penultimate layer are calculated on the entire test set after every training iteration and saved for further analysis.

We trained the network for 10 epochs (7810 training iterations) using stochastic gradient descent (SGD) with learning rate = 0.01 and variable momentum values ranging from 0 to 0.9 in 0.1 increments. Similarly, for each training iteration *t*, we sampled randomly one class (*the training class*) out of the 10 classes available and sampled 64 images randomly from only this training class to form a training batch. After updating the weights of the network using the training batch, we calculated and saved the representations of the penultimate fully-connected layer **z**_*t*_ on the entire test set for further analysis.

We then calculated the representation drift after each training iteration *t* for each class *c* as the euclidean distance between the average representations of all examplars of this class in the test set at training iteration 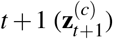 and training iteration 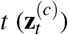 as follows:

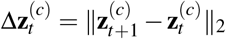

We then determined the class *c* that exhibited the highest change in representations Δ**z**^(*c*)^ after each training iteration *t*, and calculated the frequency of a class *c* that exhibited the largest change in representations when it was also the training class

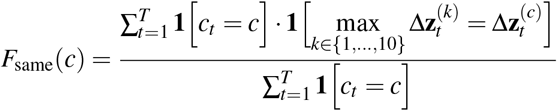

and when it was not the training class

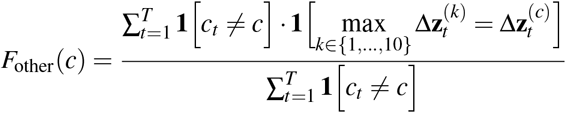

Then we computed the same-class-acceleration (SCA) ratio as 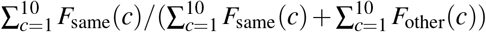. We repeated the analysis twice: once across all training iterations and once using a sliding window of 100 training iterations to track the representational drift during training.

## ACKNOWLEDGEMENT

Conceptualization: JL, MV; data analysis: JL; neural network simulations: AF, JL; mathematical analysis: JL, MV; supervision: ML; writing first draft: JL, MV; funding acquisition: MV. This work was supported by an ERC starting grant (850861) SPATEMP; DFG VI Grants (908/5-1 and 908/7-1; 505660261; 520285844; SPP LOOPS); the NWO VIDI Grant (VI.Vidi.213.124); and the Dutch Brain Interface Initiative (DBI2) of the Gravitation Program (024.005.022). We thank Giorgos Spyropoulos and Boris Sotomayor for discussions.

## SUPPLEMENTARY

**Figure S1.**
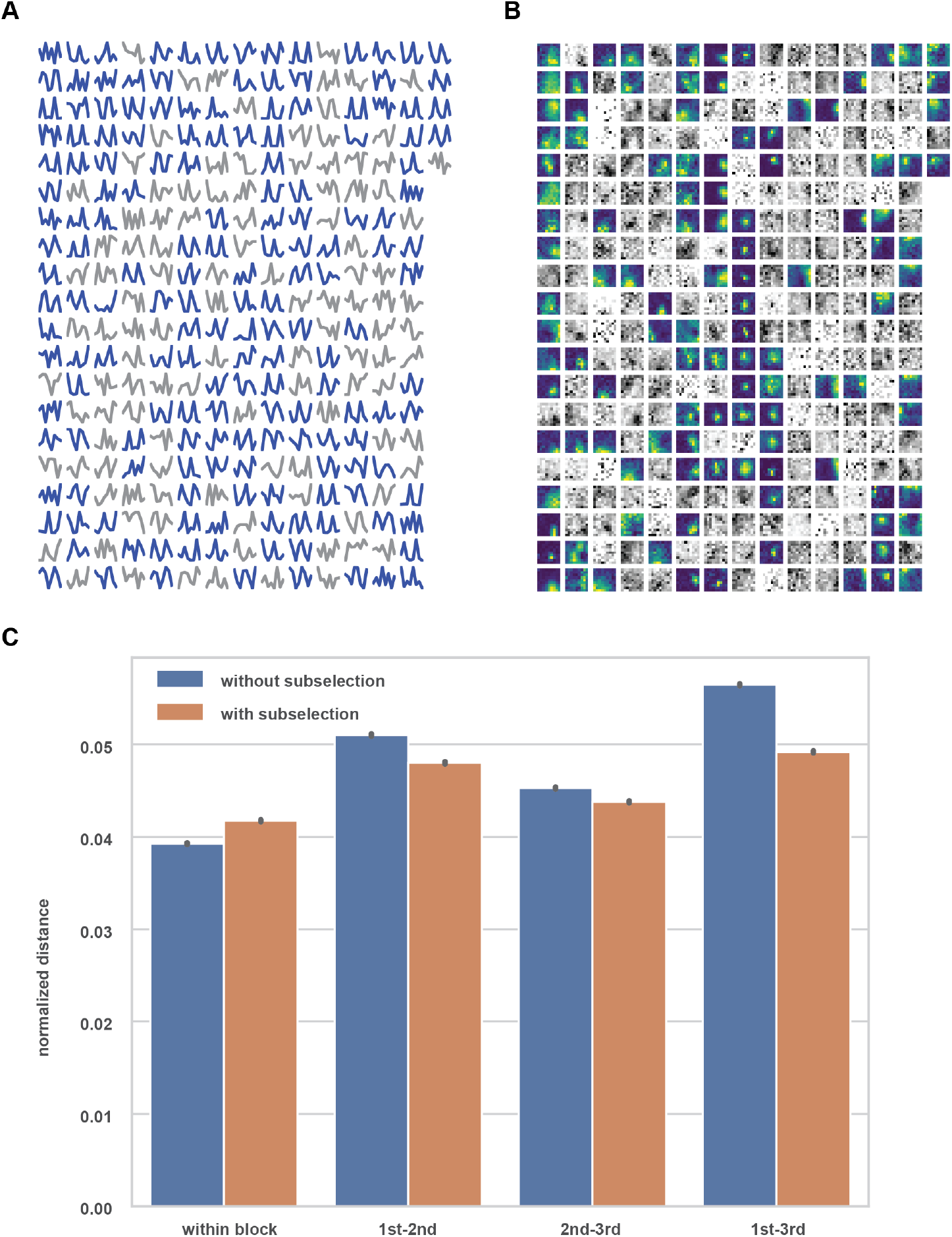
Visual neurons with only strong orientation selectivity and clear receptive field still exhibit representational drift. **A**. selecting visual neuron by its orientation tuning. **B**. selecting visual neuron by receptive field. **C**. On the low dimensional neural manifold, pairwise distances within block are significantly smaller than the distances between blocks. By selecting neurons with only strong orientation and clear receptive field, the differences shrink but still exist.

**Figure S2.**
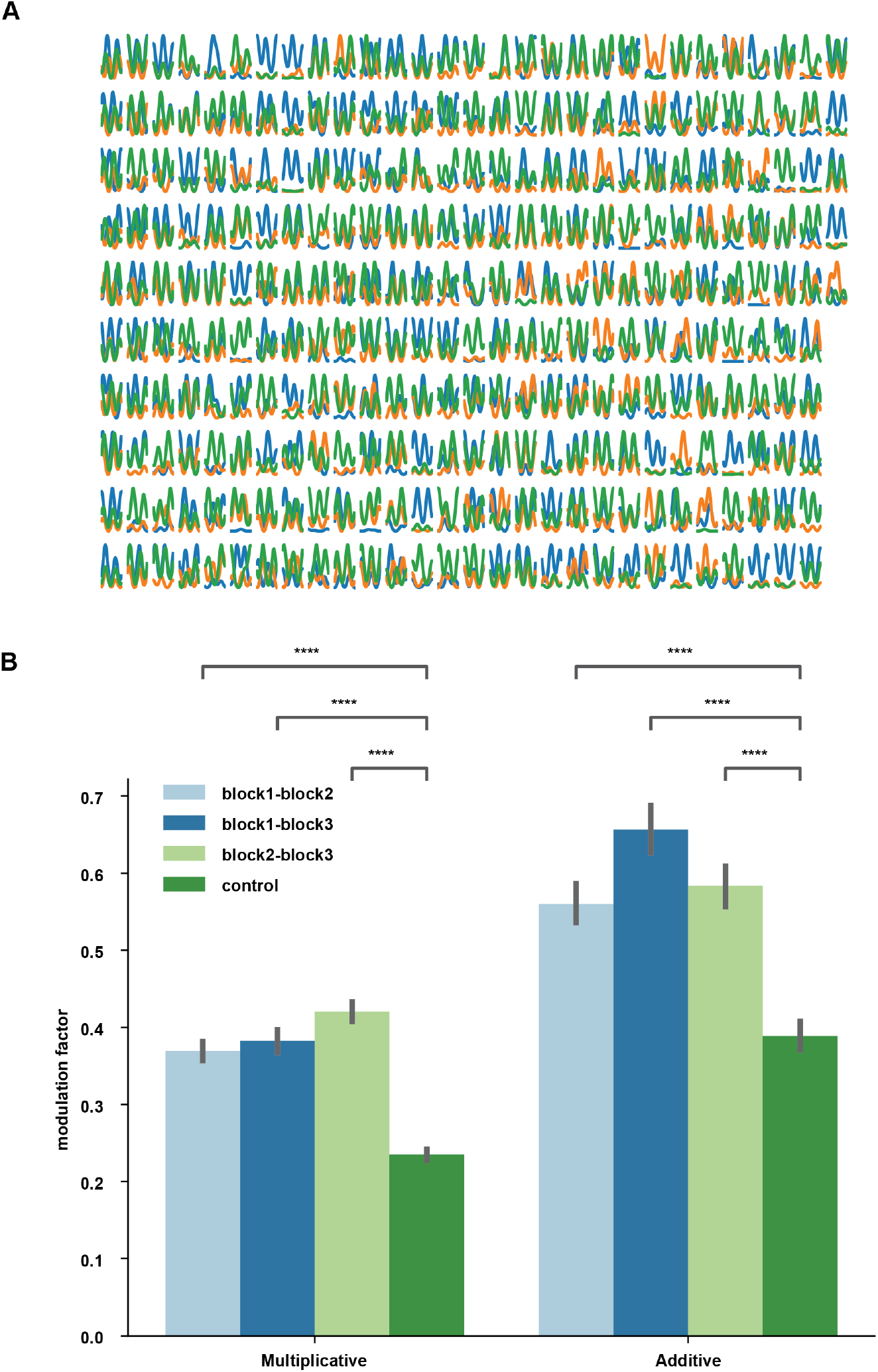
Tuning curves were modulated by block in additive and multiplicative ways. **A**. Tuning curve changes of all 289 units in example session 750749662. Tuning curves are colored according to the block identity and are fitted by Von Mises function. **B**. Across all animals, the neuronal population showed additive and multiplicative changes of tuning curves. Control AFs and MFs are computed by comparing tuning curves estimated from high-response and low-response trials.

**Figure S3.**
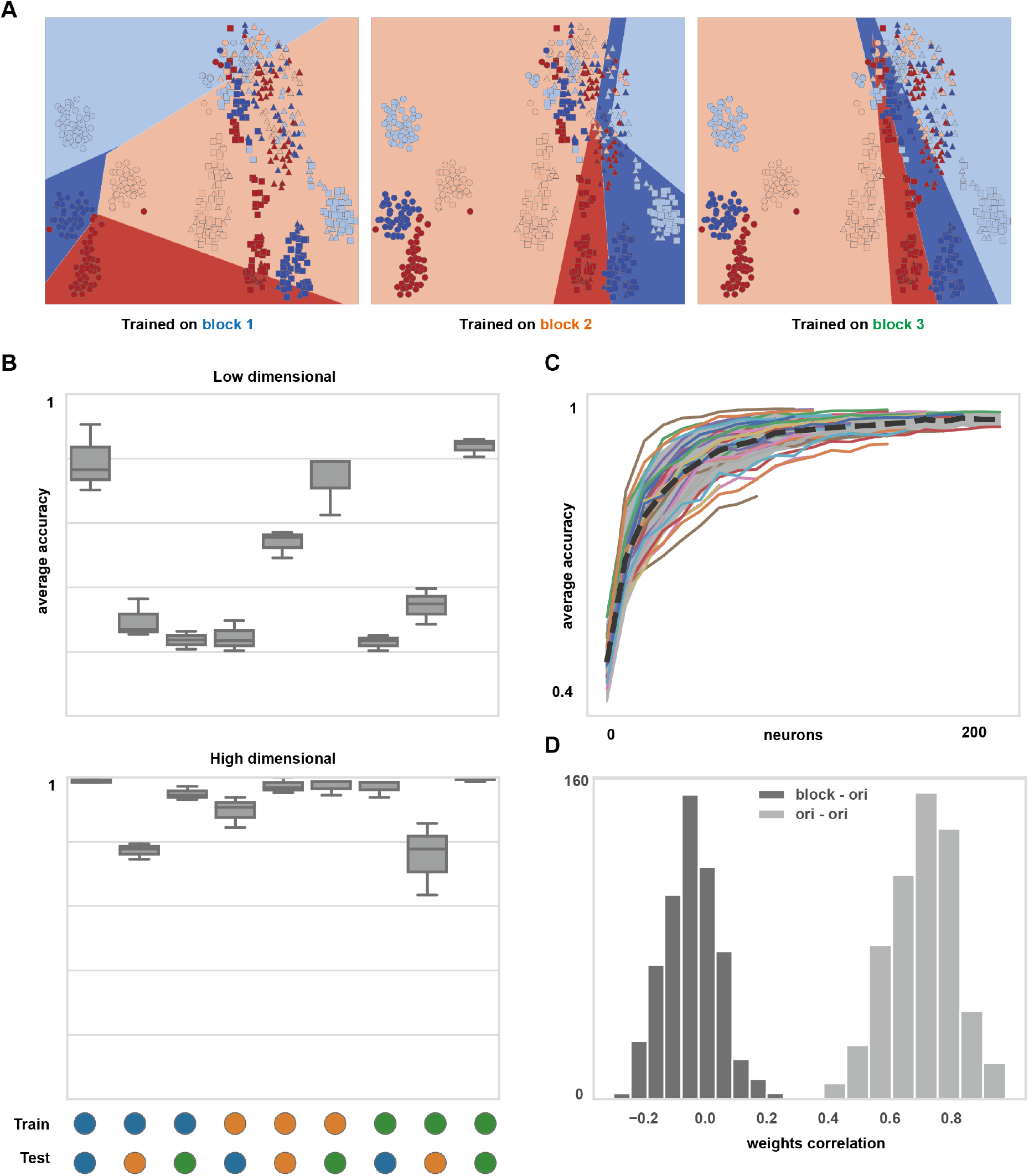
Representational drift only affects the decoding performance in low-dimensional space. **A**. In t-SNE low-dimensional representations, SVM trained on one block has poor performance on other blocks. Orientations are color labelled. Circles, triangles and squares represent the trials in the 1st, 2nd and 3rd block respectively. **B**. The cross-validated accuracy for linear classifier in low-dimensional embeddings (upper) and high-dimensional neural state space (lower). The training set and held-out test set is taken from different blocks (blue: 1st; orange: 2nd; green: 3rd) **C**. The average cross-validated accuracy for heterogeneous block train-test pairs (1st-2nd; 1st-3rd; 2nd-1st; 2nd-3rd; 3rd-1st; 3rd-2nd) increases as a function of number of neurons. Different animals were labelled in different colors and the average cross-validated accuracy of all animals was plotted in black. **D**. Distribution of correlation coefficients between weights of classifiers. Block classifiers tend to be orthogonal to orientation classifiers, while orientation classifiers of different blocks tend to be similar.

**Figure S4.**
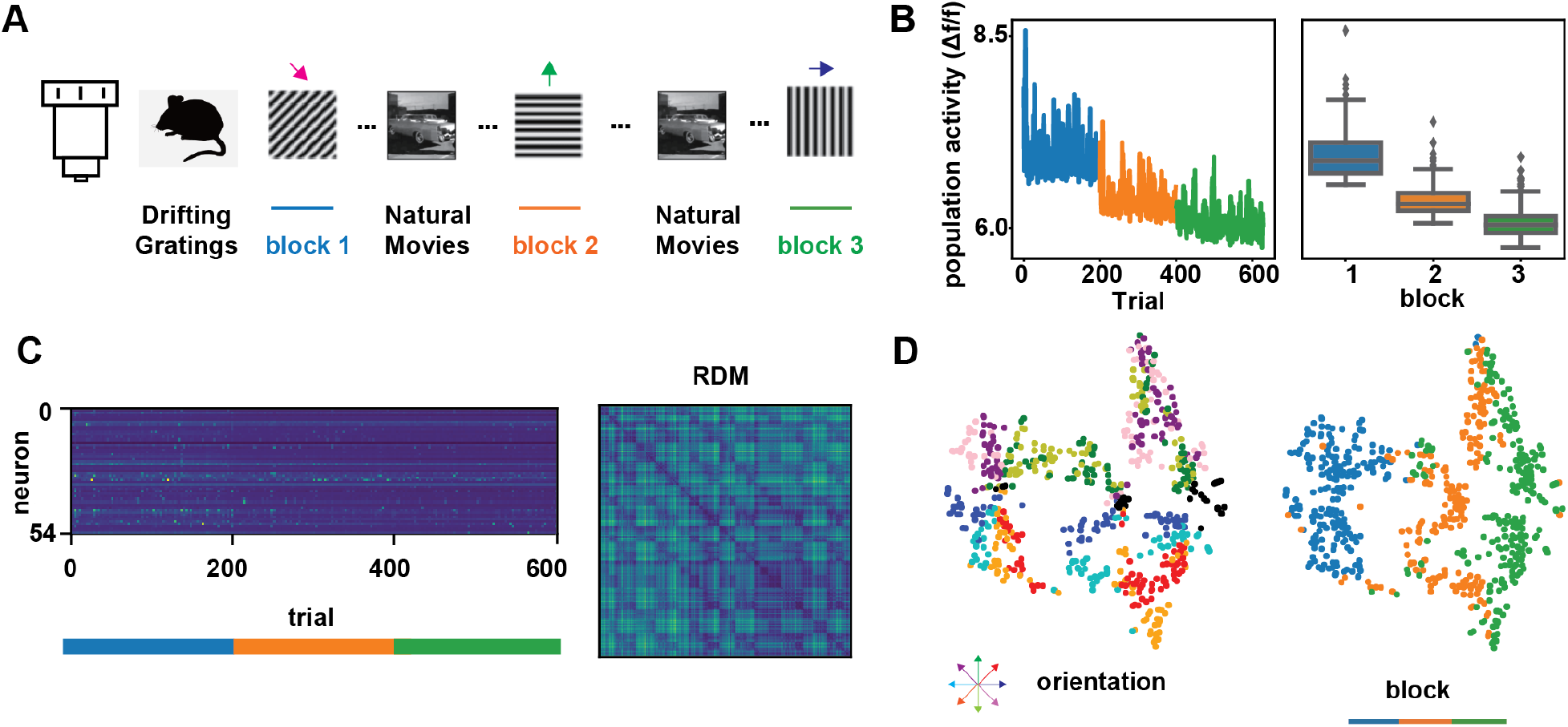
Neural representational drift causes manifold shift across blocks of drifting grating stimulus. Similar results can be observed in the calcium imaging dataset (Figure S4, de Vries et al. (2020)). We measured the responsiveness of neurons as the average Δ*F/F* value within 2 seconds after the stimulus onset considering the slower temporal dynamics of the recording methods. Similarly, the RDM exhibited orientation clustering, and the low-dimensional manifolds were separated for three different blocks (Figure S4). The comparable results from a different recording modality provided more evidence supporting that the representational drift was not merely a result of a shift of the recording device. **A-E:** An example session of 2-photon calcium imaging (511510896). **A**. Schematic illustration of calcium imaging. **B**. Overall neural response to drifting grating in each block. The neural activity was defined as the mean Δ*F/F* signal of all recorded units and all trials within 2 seconds after the stimulus onset. **C**. Population response matrix (n=54) and RDM for calcium imaging datasets. **D**. t-SNE embedding based on RDM. *upper*, labelled by drifting grating direction. The inset shows the color for each direction. *lower*, labelled by block.

**Figure S5.**
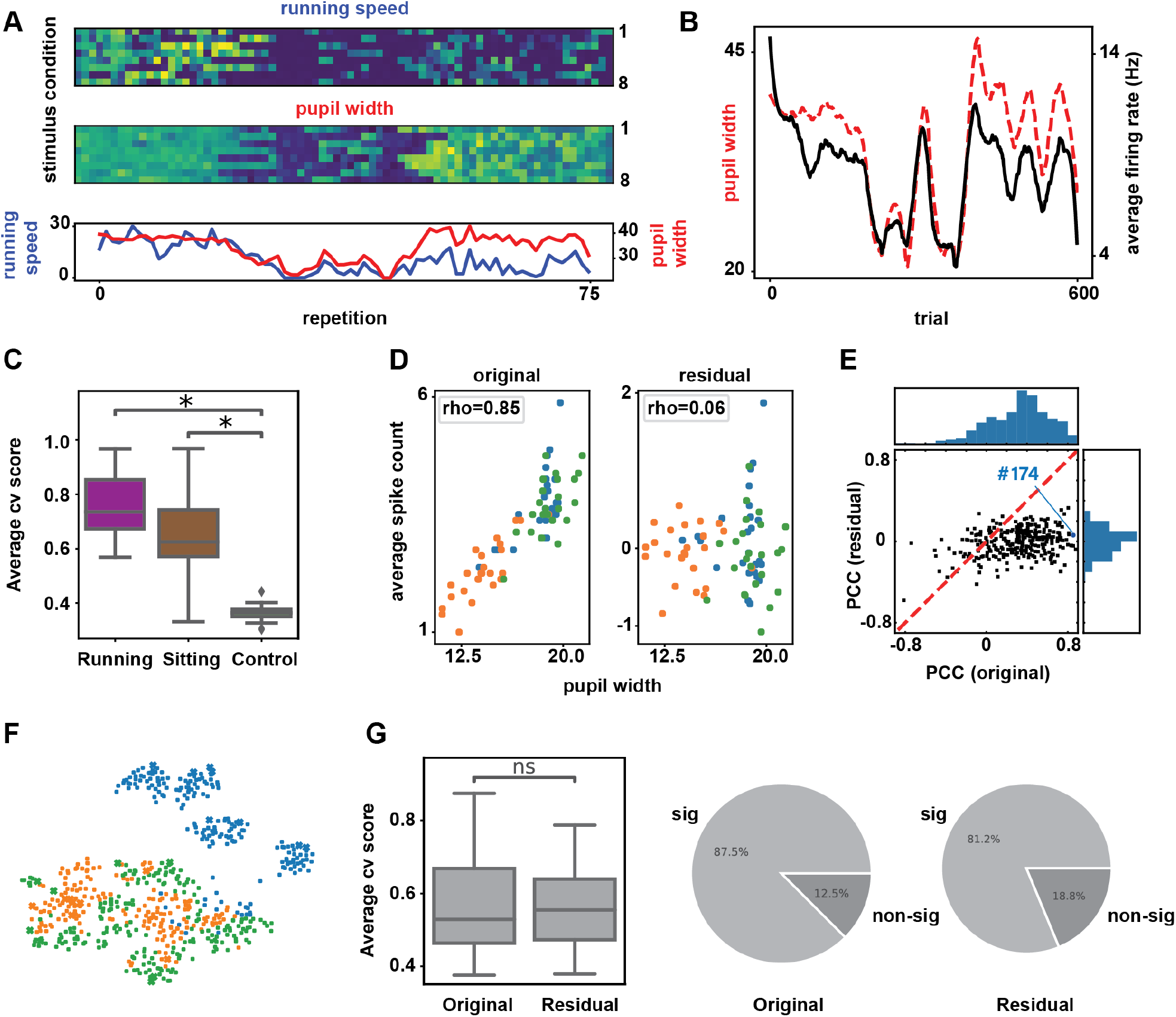
Representational drift is partially explained by animal’s behavior. **A**. The dynamics of behavioral state during the session. *upper*, average running speed of the animal for each trial. *middle*, average pupil width of the animal for each trial. *lower*, mean running speed (blue) / pupil width (red) across 8 different conditions. **B**. Correlation between pupil diameter (red) and overall mean responsiveness (black). The curves were smoothed by a Savitzky–Golay filter. **C**. Among all animals, block classifier performance on only sitting or running trials were both significantly better than chance level. **D**. The correlation between the neural response of an example neuron (#951878032) and the arousal state was broken by keeping only the residuals of a trained linear regression model. *left*, a strong correlation between the spike counts and the average pupil width; *right*, a weak correlation between the residual spike counts and the average pupil width. **E**. After keeping only the residuals of linear regression models, Pearson Correlation Coefficient (PCC) between average pupil width and average response decreased. The blue dot indicates the example neuron in panel D. **F**. Low-dimensional t-SNE embedding of residual population responses. Trials from three blocks were labeled in different colors. **G**. Block classifier performed equally well on embeddings of the original dataset and the residual dataset. More than 80% sessions showed significant block modulation even when the linear influence of behavior is removed.

**Figure S6.**
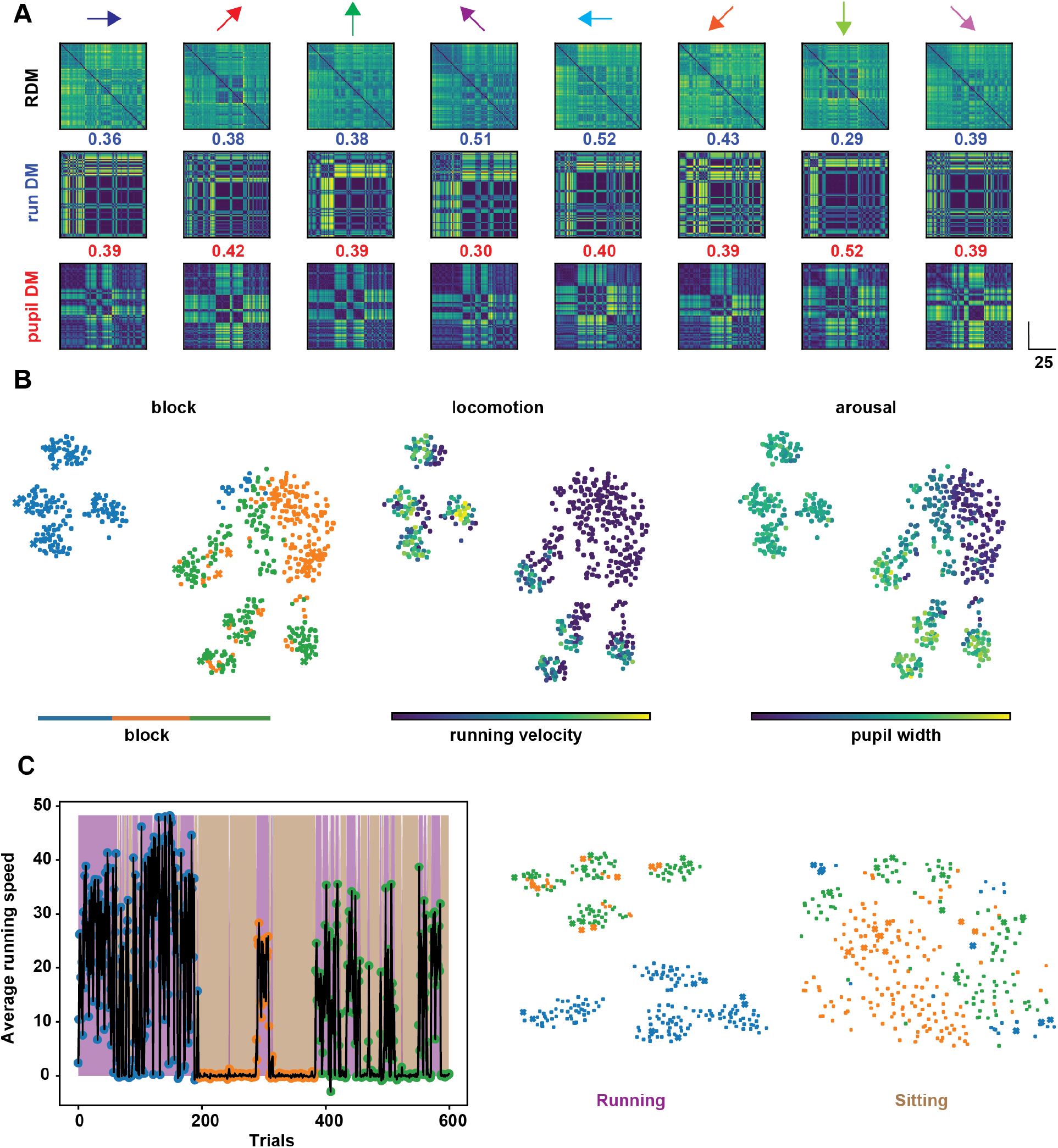
Relationship between representational drift and behavioral dynamics. Example session of 750749662. **A**. Neural representational dissimilarity matrices (RDM) showed similar patterns as behavioral state changes. *upper*, RDM for trials of each orientation; *middle*, running speed dissimilarity matrix for trials of each orientation. The average correlation between RDM and the running speed dissimilarity matrix is 0.405. *lower*, pupil width dissimilarity matrix for trials of each orientation. The average correlation between RDM and the running speed dissimilarity matrix is 0.40. The correlation between the two matrices was measured as the Pearson correlation between all off-diagonal entries. **B**. Low-dimensional neural embedding was color labelled by block, running speed, and pupil width respectively. **C**. *left*, Average running speed of the animal during the presentation of drifting grating stimulus. Trials were colored according to block. Hidden Markov Model was used to classify the animal’s behavioral state. Sitting trials were indicated as color brown while running trials as color purple. *right*, Low-dimensional neural t-SNE embedding of sitting/running trials.

**Figure S7.**
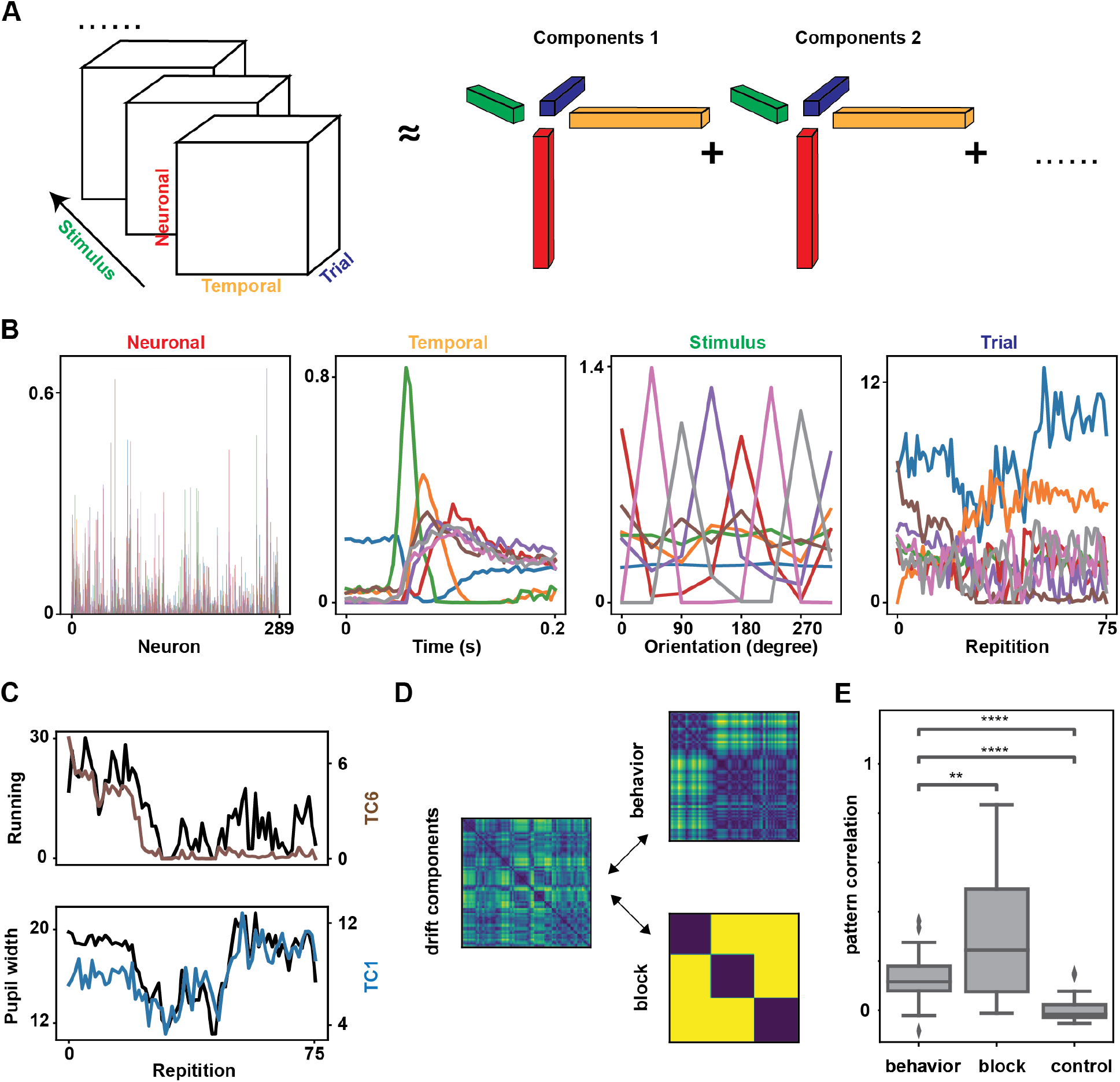
Tensor component analysis finds trial component for the representational drift. **A**. Schematic illustration of TCA algorithm on neural response as a 4D tensor with dimensions *N* × *T* × *S* × *K*. TCA reconstructs the data as a sum of rank-one tensors. Each rank-one tensor is an outer product of four components: neuronal component (red), temporal component (yellow), stimulus component (green), and trial component (blue). The set of trial components that TCA extracts describe how activity changes across 75 repetitions. **B**. TCA decomposed the 4D tensor neural response into 8 components. **C**. Trial components (TC) that were highly correlated with locomotion/arousal state dynamics. **D**. The correlation dissimilarity matrix based on the selected drift components (left). Behavior/block dissimilarity matrix (right). **E**. Pattern correlation between drift component dissimilarity matrices and behavior/block dissimilarity matrices across all sessions.

**Figure S8.**
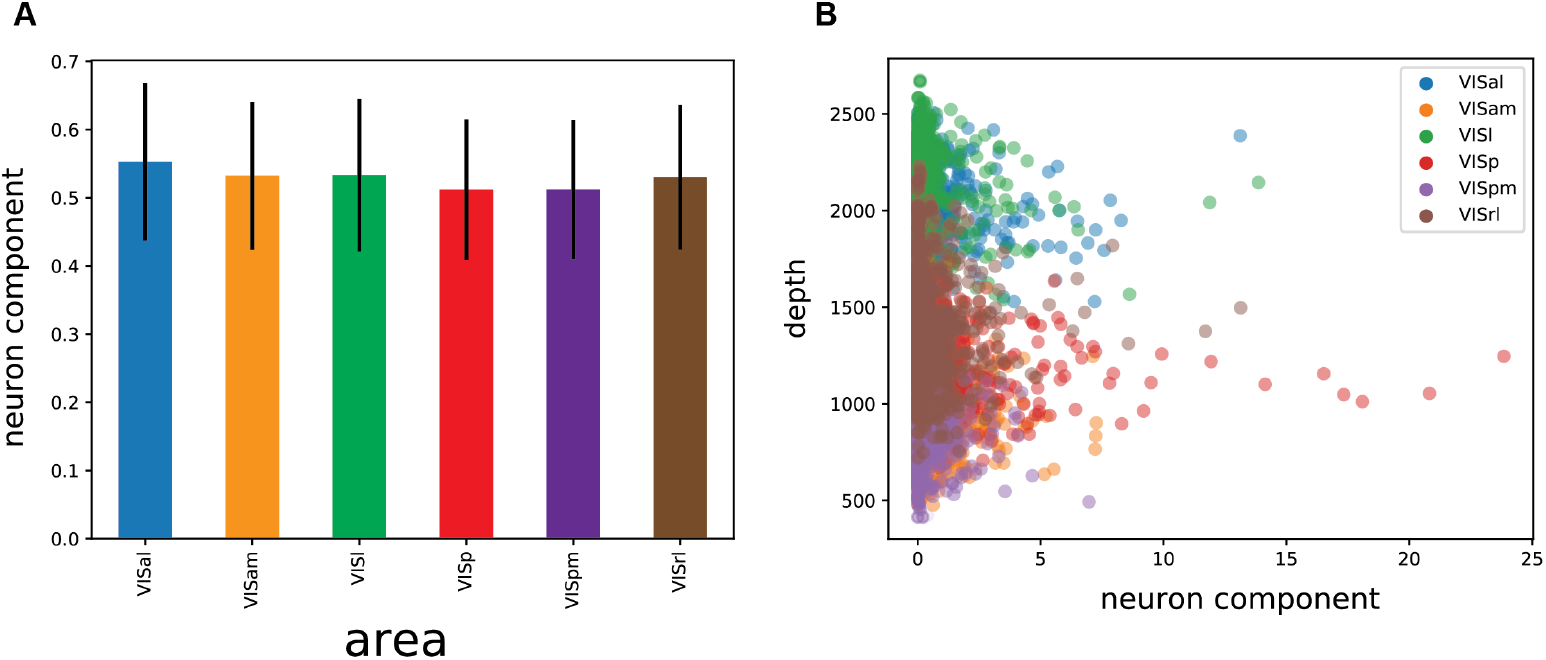
Neuronal component analysis suggests that the representational drift was distributive among the visual neuronal ensemble. **A**. Average neuronal component loadings of units in different visual areas. **B**. No correlation between the neuronal component loading and its dorsal-ventral depth.

## REFERENCES

Aitken, K., Garrett, M., Olsen, S., and Mihalas, S. (2022). The geometry of representational drift in natural and artificial neural networks. PLOS Computational Biology, 18(11):e1010716.

Aizenbud, I., Audette, N., Auksztulewicz, R., Basiński, K., Bastos, A. M., Berry, M., Canales-Johnson, A., Choi, H., Clopath, C., Cohen, U., et al. (2025). Neural mechanisms of predictive processing: a collaborative community experiment through the openscope program. arXiv preprint arXiv:2504.09614.

Arandia-Romero, I., Tanabe, S., Drugowitsch, J., Kohn, A., and Moreno-Bote, R. (2016). Multiplicative and additive modulation of neuronal tuning with population activity affects encoded information. Neuron, 89(6):1305– 1316.

Aschauer, D. F., Eppler, J.-B., Ewig, L., Chambers, A. R., Pokorny, C., Kaschube, M., and Rumpel, S. (2019). A basis set of elementary operations captures recombination of neocortical cell assemblies during basal conditions and learning. SSRN Electronic Journal.

Attardo, A., Fitzgerald, J. E., and Schnitzer, M. J. (2015). Impermanence of dendritic spines in live adult ca1 hippocampus. Nature, 523(7562):592–596.

Avitan, L. and Stringer, C. (2022). Not so spontaneous: Multi-dimensional representations of behaviors and context in sensory areas. Neuron.

Bauer, J., Lewin, U., Herbert, E., Gjorgjieva, J., Schoonover, C., Fink, A., Rose, T., Bonhoeffer, T., and Hübener, M. (2023). Sensory experience steers representational drift in mouse visual cortex. bioRxiv, pages 2023–09.

Bimbard, C., Sit, T. P., Lebedeva, A., Reddy, C. B., Harris, K. D., and Carandini, M. (2023). Behavioral origin of sound-evoked activity in mouse visual cortex. Nature Neuroscience, pages 1–8.

Brunet, N. M., Bosman, C. A., Vinck, M., Roberts, M., Oostenveld, R., Desimone, R., De Weerd, P., and Fries, P. (2014). Stimulus repetition modulates gamma-band synchronization in primate visual cortex. Proceedings of the National Academy of Sciences, 111(9):3626–3631.

Buzsáki, G. and Tingley, D. (2018). Space and time: the hippocampus as a sequence generator. Trends in cognitive sciences, 22(10):853–869.

Chambers, A. R. and Rumpel, S. (2017). A stable brain from unstable components: Emerging concepts and implications for neural computation. Neuroscience, 357:172–184.

Chen, J. L., Margolis, D. J., Stankov, A., Sumanovski, L. T., Schneider, B. L., and Helmchen, F. (2015). Pathway-specific reorganization of projection neurons in somatosensory cortex during learning. Nature Neuroscience, 18(8):1101–1108.

Cowley, B. R., Snyder, A. C., Acar, K., Williamson, R. C., Byron, M. Y., and Smith, M. A. (2020). Slow drift of neural activity as a signature of impulsivity in macaque visual and prefrontal cortex. bioRxiv.

de Vries, S. E., Lecoq, J. A., Buice, M. A., Groblewski, P. A., Ocker, G. K., Oliver, M., Feng, D., Cain, N., Ledochowitsch, P., Millman, D., et al. (2020). A largescale standardized physiological survey reveals functional organization of the mouse visual cortex. Nature Neuroscience, 23(1):138–151.

Deitch, D., Rubin, A., and Ziv, Y. (2021). Representational drift in the mouse visual cortex. Current Biology, 31(19):4327–4339.

Delamare, G., Zaki, Y., Cai, D. J., and Clopath, C. (2024). Drift of neural ensembles driven by slow fluctuations of intrinsic excitability. Elife, 12:RP88053.

Devalle, F., Zou, L., Cecchini, G., and Roxin, A. (2025). Representational drift as the consequence of ongoing memory storage. Scientific Reports, 15(1):27746.

Driscoll, L. N., Duncker, L., and Harvey, C. D. (2022). Representational drift: Emerging theories for continual learning and experimental future directions. Current opinion in neurobiology, 76:102609.

Dvorkin, R. and Ziv, N. E. (2016). Relative contributions of specific activity histories and spontaneous processes to size remodeling of glutamatergic synapses. PLoS biology, 14(10):e1002572.

Fahle, M. (2005). Perceptual learning: Specificity versus generalization. Current Opinion in Neurobiology, 15(2):154–160.

Graf, A. B., Kohn, A., Jazayeri, M., and Movshon, J. A. (2011). Decoding the activity of neuronal populations in macaque primary visual cortex. Nature neuroscience, 14(2):239–245.

Hennig, J. A., Oby, E. R., Golub, M. D., Bahureksa, L. A., Sadtler, P. T., Quick, K. M., Ryu, S. I., Tyler-Kabara, E. C., Batista, A. P., Chase, S. M., et al. (2021). Learning is shaped by abrupt changes in neural engagement. Nature Neuroscience, 24(5):727–736.

Hofer, S. B., Ko, H., Pichler, B., Vogelstein, J., Ros, H., Zeng, H., Lein, E., Lesica, N. A., and Mrsic-Flogel, T. D. (2011). Differential connectivity and response dynamics of excitatory and inhibitory neurons in visual cortex. Nature neuroscience, 14(8):1045–1052.

Keinath, A. T., Mosser, C.-A., and Brandon, M. P. (2022). The representation of context in mouse hippocampus is preserved despite neural drift. Nature communications, 13(1):2415.

Krakauer, J. W., Ghilardi, M.-F., and Ghez, C. (2006). Generalization of motor learning depends on the history of prior action. PLoS Biology, 4(10):e316.

Lange, G., van Leeuwen, J., van der Smagt, M., and Gielen, S. C. A. M. (2018). Limited transfer of visual skill in orientation discrimination across locations. Vision Research, 152:36–44.

Loewenstein, Y., Yanover, U., and Rumpel, S. (2015). Predicting the dynamics of network connectivity in the neocortex. Journal of Neuroscience, 35(36):12535–12544.

Low, I. I., Williams, A. H., Campbell, M. G., Linderman, S. W., and Giocomo, L. M. (2020). Dynamic and reversible remapping of network representations in an unchanging environment. bioRxiv.

McGinley, M. J., David, S. V., and McCormick, D. A. (2015). Cortical membrane potential signature of optimal states for sensory signal detection. Neuron, 87(1):179–192.

Micou, C. and O’Leary, T. (2023). Representational drift as a window into neural and behavioural plasticity. Current opinion in neurobiology, 81:102746.

Mongillo, G., Rumpel, S., and Loewenstein, Y. (2017). Intrinsic volatility of synaptic connections—a challenge to the synaptic trace theory of memory. Current opinion in neurobiology, 46:7–13.

Nayebi, A., Kong, N. C., Zhuang, C., Gardner, J. L., Norcia, A. M., and Yamins, D. L. (2023). Mouse visual cortex as a limited resource system that self-learns an ecologically-general representation. PLOS Computational Biology, 19(10):e1011506.

Niell, C. M. and Stryker, M. P. (2010). Modulation of visual responses by behavioral state in mouse visual cortex. Neuron, 65(4):472–479.

Pachitariu, M., Stringer, C., Dipoppa, M., Schröder, S., Rossi, L. F., Dalgleish, H., Carandini, M., and Harris, K. D. (2017). Suite2p: beyond 10,000 neurons with standard two-photon microscopy. BioRxiv.

Peter, A., Stauch, B. J., Shapcott, K., Kouroupaki, K., Schmiedt, J. T., Klein, L., Klon-Lipok, J., Dowdall, J. R., Schölvinck, M. L., Vinck, M., et al. (2021). Stimulus-specific plasticity of macaque v1 spike rates and gamma. Cell reports, 37(10).

Psarou, E., Parto-Dezfouli, M., Grothe, I., Peter, A., Roese, R., and Fries, P. (2025). Repetition-related gamma plasticity in macaque v1 and v2 is highly stimulus specific and robust to stimulus set size. bioRxiv, pages 2025–06.

Qin, S., Farashahi, S., Lipshutz, D., Sengupta, A. M., Chklovskii, D. B., and Pehlevan, C. (2023). Coordinated drift of receptive fields in hebbian/anti-hebbian network models during noisy representation learning. Nature Neuroscience, pages 1–11.

Raman, D. V. and O’Leary, T. (2021). Optimal plasticity for memory maintenance during ongoing synaptic change. elife, 10:e62912.

Rokni, U., Richardson, A. G., Bizzi, E., and Seung, H. S. (2007). Motor learning with unstable neural representations. Neuron, 54(4):653–666.

Rule, M. E., Loback, A. R., Raman, D. V., Driscoll, L. N., Harvey, C. D., and O’Leary, T. (2020). Stable task information from an unstable neural population. Elife, 9:e51121.

Rule, M. E. and O’Leary, T. (2022). Self-healing codes: How stable neural populations can track continually reconfiguring neural representations. Proceedings of the National Academy of Sciences, 119(7):e2106692119.

Rule, M. E., O’Leary, T., and Harvey, C. D. (2019). Causes and consequences of representational drift. Current opinion in neurobiology, 58:141–147.

Sadeh, S. and Clopath, C. (2022). Contribution of behavioural variability to representational drift. Elife, 11:e77907.

Schoonover, C. E., Ohashi, S. N., Axel, R., and Fink, A. J. (2021). Representational drift in primary olfactory cortex. Nature, pages 1–6.

Siegle, J. H., Jia, X., Durand, S., Gale, S., Bennett, C., Graddis, N., Heller, G., Ramirez, T. K., Choi, H., Luviano, J. A., et al. (2021). Survey of spiking in the mouse visual system reveals functional hierarchy. Nature, 592(7852):86–92.

Singh, A., Peyrache, A., and Humphries, M. D. (2019). Medial prefrontal cortex population activity is plastic irrespective of learning. Journal of Neuroscience, 39(18):3470–3483.

Sotomayor-Gomez, B., Battaglia, F. P., and Vinck, M. (2023). Differential population coding of natural movies through spike counts and temporal sequences. bioRxiv, pages 2023–06.

Stringer, C., Pachitariu, M., Steinmetz, N., Reddy, C. B., Carandini, M., and Harris, K. D. (2019). Spontaneous behaviors drive multidimensional, brainwide activity. Science, 364(6437).

Swindale, N. V. (1998). Orientation tuning curves: empirical description and estimation of parameters. Biological cybernetics, 78(1):45–56.

Tang, W., Chang, H., Liu, C., Perez-Hernandez, S., Zheng, W. Y., Park, J., Oliva, A., and Fernandez-Ruiz, A. (2025). A hippocampal population code for rapid generalization. bioRxiv, pages 2025–03.

Uran, C., Peter, A., Lazar, A., Barnes, W., Klon-Lipok, J., Shapcott, K. A., Roese, R., Fries, P., Singer, W., and Vinck, M. (2022). Predictive coding of natural images by v1 firing rates and rhythmic synchronization. Neuron, 110(7):1240–1257.

Van der Maaten, L. and Hinton, G. (2008). Visualizing data using t-sne. Journal of machine learning research, 9(11).

Vinck, M., Batista-Brito, R., Knoblich, U., and Cardin, J. A. (2015). Arousal and locomotion make distinct contributions to cortical activity patterns and visual encoding. Neuron, 86(3):740–754.

Williams, A. H., Kim, T. H., Wang, F., Vyas, S., Ryu, S. I., Shenoy, K. V., Schnitzer, M., Kolda, T. G., and Ganguli, S. (2018). Unsupervised discovery of demixed, low-dimensional neural dynamics across multiple timescales through tensor component analysis. Neuron, 98(6):1099–1115.

Zhong, L., Baptista, S., Gattoni, R., Arnold, J., Flickinger, D., Stringer, C., and Pachitariu, M. (2025). Unsupervised pretraining in biological neural networks. Nature, pages 1–8.

Ziv, Y., Burns, L. D., Cocker, E. D., Hamel, E. O., Ghosh, K. K., Kitch, L. J., El Gamal, A., and Schnitzer, M. J. (2013). Long-term dynamics of ca1 hippocampal place codes. Nature neuroscience, 16(3):264–266.

